# Single-turnover kinetic analysis of non-LTR retrotransposition defines the mechanism and rate constants governing each step

**DOI:** 10.1101/2024.08.28.610141

**Authors:** Tyler L. Dangerfield, Jun Zhou, Thomas Eickbush, Kenneth A. Johnson

## Abstract

Site specific retrotransposon-mediated gene therapy has the potential to revolutionize medicine by allowing insertion of large gene cargos. Despite decades of effort, the reaction sequence remains to be fully elucidated limiting the ability to engineer improved activity for gene insertion. Here we provide a kinetic/mechanistic framework for R2 non-LTR retrotransposition. Single turnover measurements and global data fitting defined the rate constants for each step in the pathway involving 1^st^-strand DNA cleavage to provide a DNA primer, reverse transcription to copy the RNA, 2^nd^-strand DNA cleavage to provide the second primer, and 2^nd^-strand synthesis to make duplex cDNA. Sequence analysis of the cDNA confirms accurate replication of the 1400 nt RNA used in this study. This represents the only complete analysis of the reaction sequence and first observation of 2^nd^-strand synthesis *in vitro*. We provide a kinetic framework to understand non-LTR retrotransposition, which provide a basis to engineer improved activity.

## Introduction

Non-long-terminal repeat (non-LTR) retrotransposons (also called <u>L</u>ong Interspersed <u>N</u>uclear <u>E</u>lements, LINEs) are abundant in eukaryotic genomes. R2 elements are a subset of LINEs that insert site-specifically into the 28S rRNA genes of their host (**Figure 1**) and are found in many animal phyla but not mammals ^11^. Unlike the non-LTR retrotransposons (L1) found in humans, R2 elements encode a single open reading frame that contains the activities required for the retrotransposition reaction; namely, zinc finger and c-myb domains for target site recognition, an endonuclease domain, and a reverse transcriptase domain (**Figure 1A**). The most well characterized R2 element is R2Bm from the silk moth, *Bombyx mori* ^2,3^. After the endonuclease nicks the DNA target, the 3’ end of the cleaved DNA primes reverse transcription using the R2 mRNA as a template in a process called <u>T</u>arget <u>P</u>rimed <u>R</u>everse <u>T</u>ranscription (TPRT)^4^. The enzyme then is postulated to nick the second DNA strand, providing a primer for 2^nd^-strand DNA synthesis which uses the 1^st^ strand TPRT product as a template to generate duplex cDNA. Highly structured RNA from two regions of the R2 transcript, the 3’ untranslated region (3’UTR) and a region encoding the N-terminal part of the R2 open reading frame have been shown to be critical for R2 insertion. These RNAs specifically bind to the enzyme via their complex secondary structures^3,5–7^. Transposable elements have evolved for their long-term survival in host genomes and not necessarily for rapid and efficient new integrations. As a result, many elements may have low activity and efficiency. To repurpose these elements as gene insertion tools, greater mechanistic understanding is requisite to engineer the enzyme for increased efficiency in human cells.

**Figure 1:**
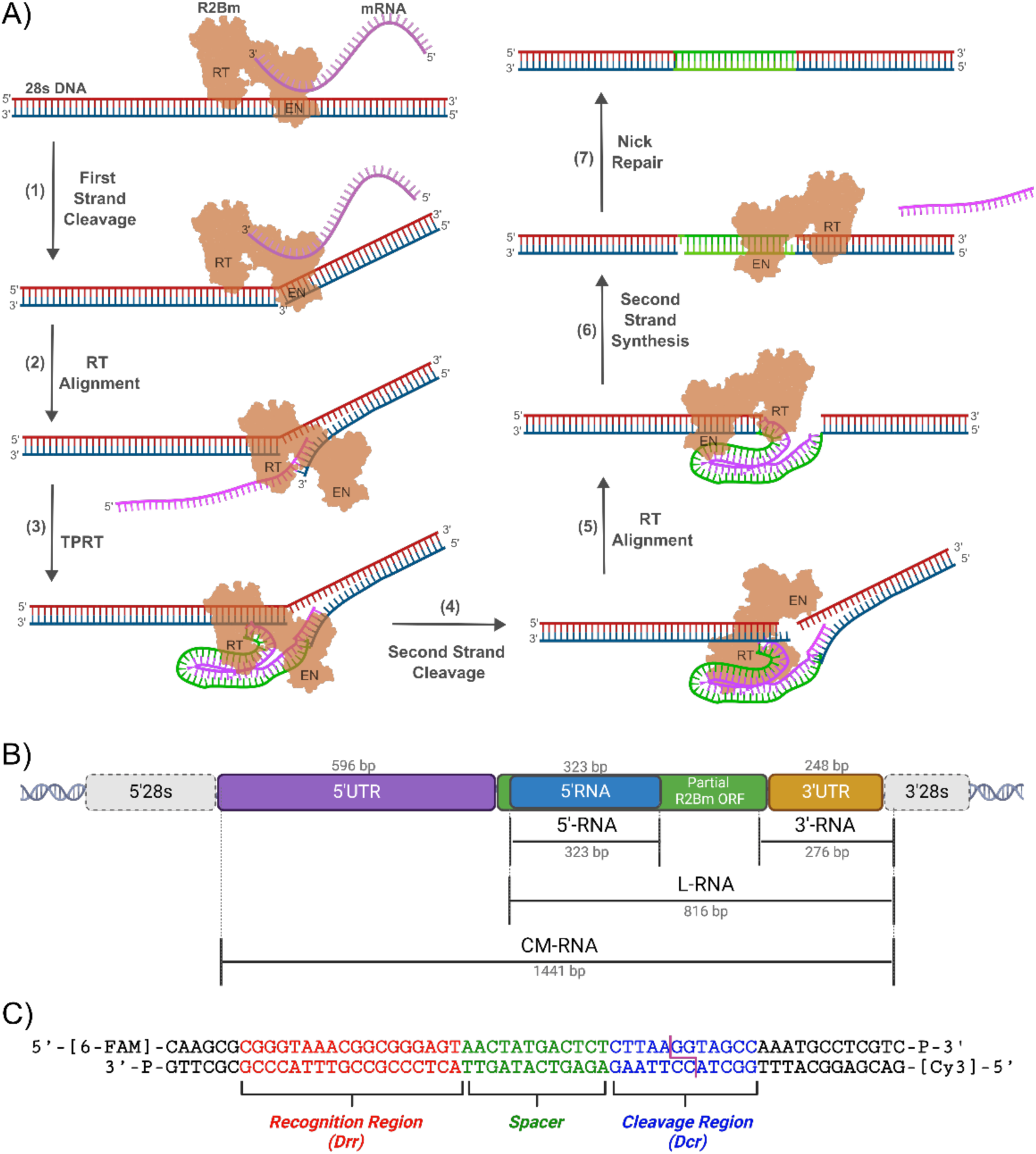
**R2Bm Mechanism Cartoon, Gene Structure, and 28S DNA Target**. <u>A) Cartoon of R2</u> <u>retrotransposition reaction pathway.</u> R2 enzyme (brown), bound to its mRNA (pink), binds the 28S DNA locus and nicks the first strand (1). This is followed by transfer of the nicked DNA from the endonuclease active site (EN) to the polymerase active site (RT) (2). TPRT then occurs using the R2 mRNA as a template and the nicked lower strand as a primer (3). The second strand is then nicked using the 5’RNA cofactor and the new 3’ end is used as a primer to copy the cDNA from the TPRT reaction (4, 5, and 6). Finally, host factors seal the nicks, leaving an R2 element integrated in the host genome (7). For simplicity in drawing, the diagram contains one R2 protein, but the reaction could involve a dimer. Only the polymerase and endonuclease domain of the protein is drawn. A third sequence specific DNA binding domain is also present in the protein but is not the subject of this report. <u>B) R2Bm gene structure and RNAs used in this study</u>. Flanking 28S regions are shown in grey, the 5’UTR is shown in purple, and the truncated R2Bm ORF is shown in green. The two regions of the elements whose RNA sequence is known to be recognized by the R2 protein correspond to the 5’RNA segment shown in blue, and the 3’UTR is shown in yellow. RNAs used in this study are diagrammed below along with their length. The truncated ORF, containing the 5’RNA region is 582 nt long while the full R2Bm ORF (not shown) is 3342 nt long. <u>C) 28s DNA target.</u> The 28s DNA target is shown, annotated with regions that R2Bm recognizes, based on findings by Liu, *et al.*^777^. A 3’ phosphate blocking group was added to both strands to prevent non-specific addition to the 3’ ends of the starting DNA substrate. Top and bottom strands are 5’ labelled with 6-FAM and Cy3, respectively.

The stepwise model previously proposed for TPRT^2^ has been illuminated by cryo-EM analysis^7,8^, but structure alone does not define the pathway or identify the rate-limiting and specificity-determining steps. Indeed, little is known about the kinetics of the overall reaction, particularly the steps following TPRT. While 2^nd^-strand cleavage and subsequent DNA-dependent 2^nd^-strand synthesis are predicted to achieve gene insertion, 2^nd^-strand cleavage efficiency is low and 2^nd^-strand synthesis has never been directly observed. Many mechanistic questions remain for this delicate process. Does the R2 enzyme alone catalyze 2^nd^-strand cleavage and synthesis or is it dependent upon repair enzymes within a cell? Does the enzyme remain bound to RNA and DNA throughout this complex multistep reaction and, if so, how does it shuttle the DNA between endonuclease and polymerase active sites? Does R2 enzyme bind as a monomer or dimer? Why are the 5’ ends of integrated reaction products seen *in vivo* so variable? What are the levels of efficiency of the various steps?

We address many of these questions and provide a complete kinetic framework for understanding the gene insertion activity of R2Bm using single turnover kinetic methods with defined minimal mRNAs and target DNA oligonucleotides. We quantify the active site enzyme concentration and DNA binding affinity in our enzyme preparation, and define the important role played by the two specific segments of RNA in the DNA cleavage and synthesis reactions. Finally, using a complete minimal RNA, containing all structured elements from the 5’UTR through 3’UTR of the R2 element, we directly observe all four sequential reactions including 2^nd^-strand cleavage and 2^nd^-strand synthesis to generate a duplex cDNA product.

## Results

### Effect of 3’RNA on 1^st^-strand cleavage

It has been previously shown that RNA, even non-specific RNA, stimulates R2 to cleave DNA at the 28S locus insertion site. However, only RNA corresponding to the 3’ end of the R2Bm element is used as a template in an integration reaction^6,9^. As an initial measure of activity, 400 nM R2 enzyme (nominal concentration) was preincubated with or without 1 µM 3’RNA (**Figure 1B**). To start the reaction, we added the 20 nM 28S dsDNA substrate (**Figure 1C**) and then stopped the reaction at various times by adding EDTA. The DNA cleavage reaction products as a function of time were quantified by capillary electrophoresis with fluorescence detection of the 5’-labeled DNA (**Figure 1C**) to get the results shown in **Figure 2A**. Cleavage was stimulated by 3’RNA with an amplitude almost 10-fold higher than the reaction without RNA, although the observed exponential decay rates were both approximately 0.6 min^−1^. Largely irreversible reactions examined with excess enzyme are expected to go to completion, but in this case, only ∼75% of the DNA (15 of 20 nM) was cleaved after 5 minutes and no further cleavage was observed after 45 min. These results suggest that a fraction of the DNA binds to the enzyme in a non-productive (unreactive) state and does not dissociate from the enzyme on the timescale of the experiment.

**Figure 2:**
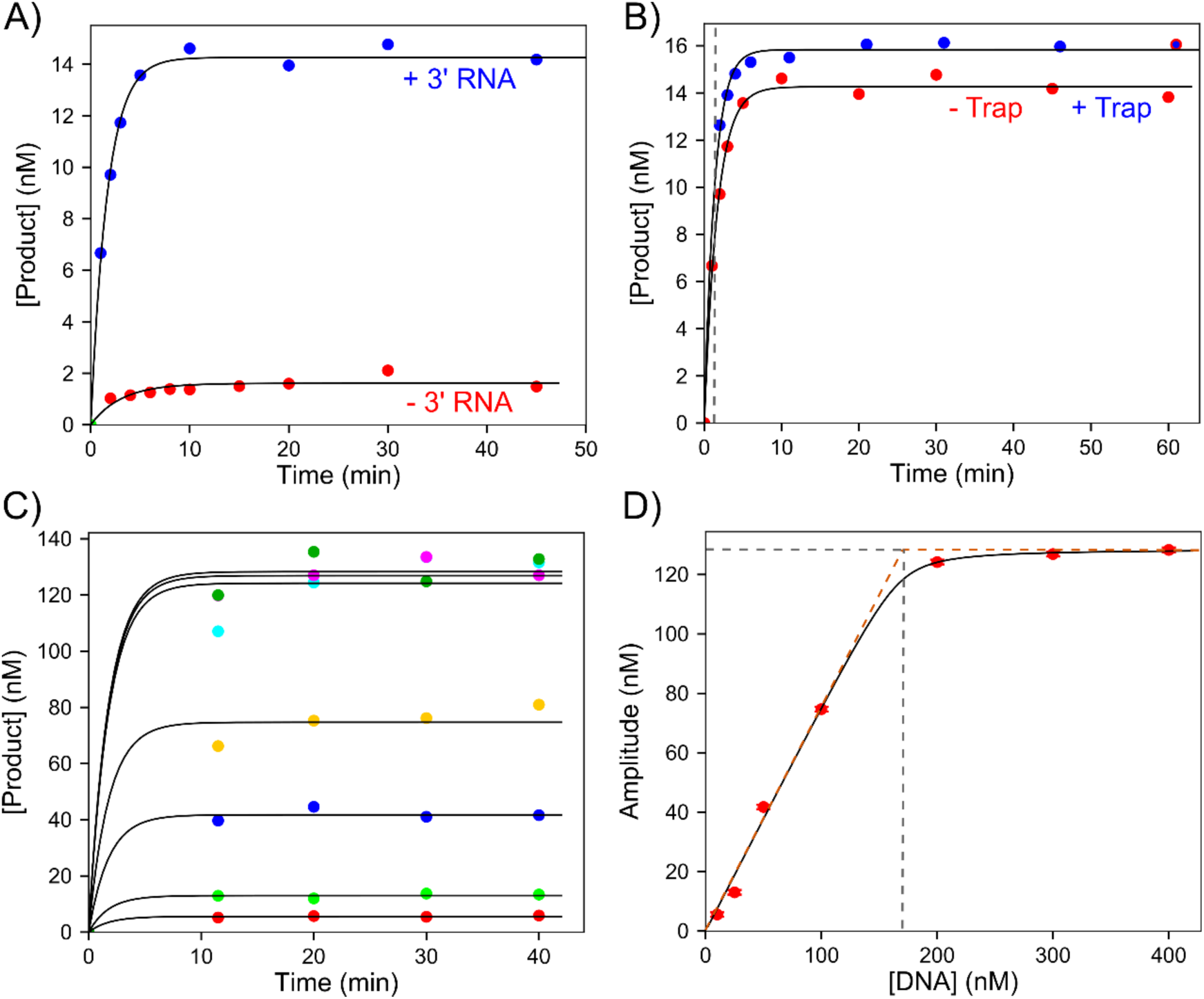
First Strand Cleavage / Active Site Titration of R2Bm. <u>A) Cleavage of the first strand with or</u> <u>without the 3’RNA</u>. A solution of 400 nM R2Bm (nominal concentration) and 0 or 1 µM 3’RNA was mixed with 20 nM DNA to start the reaction. Samples were quenched at various times with EDTA and products were resolved by capillary electrophoresis. Data shown in red were collected without RNA and data in blue were collected with RNA. Data are shown fit to single exponential functions. Both curves had an observed rate of 0.6 min^−1^ but with amplitudes of 1.6 nM and 14 nM without and with RNA, respectively. <u>B) First strand</u> <u>cleavage trapping experiment.</u> A solution of 400 nM R2Bm (concentration by absorbance) and 1 µM 3’RNA was mixed with 40 nM DNA to start the reaction. After 1 minute the reaction was mixed 1:1 with unlabeled DNA trap at a final concentration of 1 µM. The amplitude without the trap was 14 nM while the amplitude with the trap was 16 nM and both occurred with observed rates of 0.6 min^−1^. The dashed line shows the time point at which the trap was added. <u>C) Active site titration of R2Bm.</u> A solution of 400 nM R2Bm (concentration by absorbance) and 1 µM 3’RNA was mixed with varying concentrations of DNA from 10 – 400 nM and samples were quenched with EDTA at various times. Data shown in different colors represent different DNA concentrations and the black lines through the data are best fits to single exponential functions. <u>D) Amplitude versus concentration for active site titration in C.</u> The amplitudes from the single exponential fits in (C) are shown plotted as a function of DNA concentration. The black line is the best fit of the data to a quadratic equation giving a maximum amplitude of 130 ± 4 nM, a *K_d_* ⩽ 4 nm, and an enzyme concentration of 170 ± 10 nM representing 43% active enzyme relative to the nominal concentration estimated by absorbance measurements. Orange dashed lines are shown for infinitely tight binding and the grey dashed lines give the amplitude and DNA concentration at this saturation point, defining the key parameters from the fit to the quadratic equation.

In **Figure 2B**, we probed the stability of the enzyme-DNA complex by adding an excess of unlabeled DNA soon after the reaction was started. The reaction was initiated by mixing a pre-formed R2 enzyme-3’RNA complex with labelled DNA, and after 1 minute a 40-fold excess of unlabeled DNA was added to trap any free enzyme that dissociated from the labelled DNA. Any product formed after 1 minute must arise from DNA that bound to the enzyme within 1 minute and did not dissociate on the timescale of the measurement. The reaction with the trap had a similar cleavage amplitude compared to the reaction without the trap, demonstrating that once bound to the target DNA the enzyme did not dissociate before cleaving the target. These data also indicate that DNA binding must be faster than DNA cleavage so that a fast, largely irreversible DNA binding and recognition step was followed by a slower cleavage reaction (0.6 min^−1^ = 0.01 s^−1^). The slightly higher amplitude in the presence of excess unlabeled DNA (i.e. the trap) may imply that the excess DNA stimulated a slightly higher fraction of the enzyme-bound DNA to be in the active conformation.

### Active Site Titration of R2 During 1^st^-strand Cleavage

In any protein preparation there is likely to be a fraction of inactive enzyme and/or systematic errors in protein determination methods. Therefore, an important standard of any quantitative kinetic study is an accurate quantification of the active enzyme concentration relative to the nominal enzyme concentration estimated by other methods. Our method depends on having a chemistry step that is faster than the rate of substrate and product dissociation, as was demonstrated for R2 in **Figure 2B**. For active site titration, we pre-formed a complex of the R2 enzyme and 3’RNA at a fixed concentration and then measured 1^st^-strand cleavage as a function of time at various DNA concentrations (**Figure 2C**). At all DNA concentrations the reaction amplitude reached the endpoint within the first 10 min. Plotting reaction amplitude as a function of total DNA concentration (**Figure 2D**) gave a curve that best fits a quadratic function giving a *K_d_* estimate of less than 4 nM for the enzyme-DNA complex, consistent with observations in prior studies^3,10^. The active site titration (**Figure 2D**) shows that 170 nM DNA is required to saturate the amplitude of the 1^st^-strand cleavage reaction, thereby defining the active enzyme concentration. Compared to the 400 nM nominal concentration defined by absorbance measurements, we estimate ∼43% of the enzyme is able to bind the DNA target. This calculation is based on the assumption that the enzyme binds as a monomer. Of the 170 nM DNA bound to the enzyme only 130 nM product was formed (y-axis) suggesting, as does the experiment in **Figure 2A**, that ∼25% of bound DNA is in a non-productive (inactive) state. This is not unusual in that a small fraction of non-productive DNA binding was also seen in studies on CRISPR Cas9 endonuclease^111111^. A quantitative model that accounts for the rate and amplitude data is shown in **Figure 4E**, including results from additional experiments.

### Second strand cleavage

Previous studies have shown that a region of mRNA encoding the N-terminal end of the R2 ORF is required for efficient 2^nd^-strand DNA cleavage^6,7^. We therefore investigated the kinetics of 2^nd^-strand cleavage in the presence of this RNA segment which we refer to as 5’RNA (**Figure 1B**, **Table 3**). As in the experiments just described, R2 enzyme and 5’RNA were preincubated, then mixed with the DNA substrate to start the reaction. Both strands of the DNA were cleaved in this experiment and the amount of cleavage for each strand as a function of time is shown in **Figure 3A**. First-strand cleavage stimulated by 5’RNA was similar in amplitude as seen with 3’RNA but at a slightly slower rate. Second strand cleavage in the presence of 5’RNA was observed with a similar amplitude as seen with 1^st^ stand cleavage but at a five-fold slower rate. There was no noticeable lag in the 2^nd^ strand cleavage as might be expected if the two events were sequential. We next performed a trapping experiment with a large excess of unlabeled DNA added one minute after starting the reaction (**Figure 3B**). Unlike the result seen with 3’RNA (**Figure 2B**), no further 1^st^ strand cleavage occurred after the addition of the trap suggesting that when the reaction was stimulated by 5’RNA the DNA can dissociate from the enzyme faster than 1^st^ strand cleavage. Surprisingly 2^nd^-strand cleavage continued after the addition of the trap until the amplitude reached the level of 1^st^ strand cleavage. This experiment suggests that in the presence of 5’RNA, the DNA-enzyme complex is unstable, but once 1^st^ strand cleavage has occurred, the DNA is stably bound allowing 2^nd^-strand cleavage. With 5’RNA the R2 enzyme can dissociate from the DNA before 1^st^ strand cleavage but not after. In contrast, 3’RNA promotes stable binding before and after 1^st^ strand cleavage. Note that these interpretations reflect the relative rates of DNA dissociation versus cleavage. To see DNA cleavage in the presence of the trap, the dissociation rate only needs to be at least tenfold slower than cleavage.

**Figure 3:**
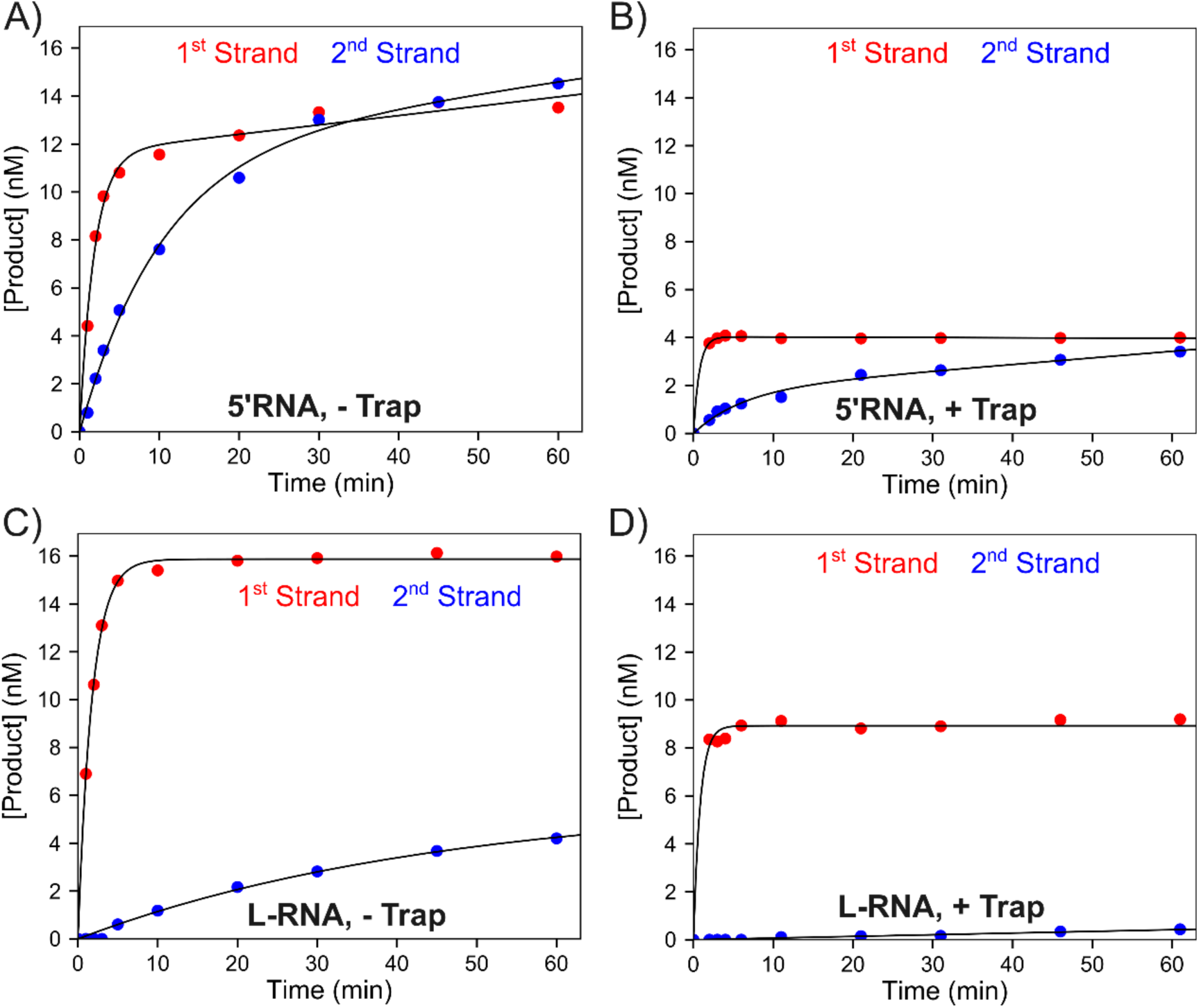
First and second strand DNA cleavage with 5’RNA and L-RNA. For each experiment in this figure, points in red show 1^st^-strand cleavage and points in blue show 2^nd^ strand cleavage. <u>A) DNA cleavage</u> <u>activity stimulated by 5’RNA.</u> A solution of 172 nM R2Bm and 1 µM 5’RNA was mixed with 20 nM DNA to start the reaction. Both data sets were fit to a single exponential burst equation. First-strand cleavage occurred with an exponential phase with an amplitude of 11.6 nM and a rate of 0.55 min^−1^ followed by a linear rate of 0.04 nM/min. Second-strand cleavage occurred with an amplitude of 11.8 nM at a rate of 0.1 min^−1^, followed by a linear rate of 0.05 nM/min. <u>B) Trapping experiment for reactions stimulated by 5’RNA.</u> A solution of 172 nM R2Bm and 1 µM 5’RNA was mixed with 40 nM DNA to start the reaction. After 1 minute, the reaction was mixed 1:1 with a solution of unlabeled DNA to give concentration of 1 µM after mixing. The amplitude of 1^st^-strand cleavage was 4 nM (rate not resolved) while the 2^nd^-strand cleavage occurred with an amplitude of 1.8 nM and rates of 0.16 min^−1^ and 0.027 min^−1^ for fast and slow phases, respectively. <u>C) DNA cleavage activity stimulated by L-RNA.</u> A solution of 172 nM R2Bm and 1 µM L-RNA was mixed with 20 nM DNA to start the reaction. Data for 1^st^ strand cleavage fit a single exponential function with an amplitude of 16 nM and an observed rate of 0.57 min^−1^. Second-strand cleavage fit a single exponential function with an amplitude of 5.8 nM and an observed rate of 0.02 min^−1^. <u>D) Trapping</u> <u>experiment for L-RNA.</u> A solution of 172 nM R2Bm and 1 µM L-RNA was mixed with 40 nM labelled DNA to start the reaction. After 1 minute, the reaction was mixed at a 1:1 ratio with unlabeled DNA at a final concentration of 1 µM. Data for 1^st^ strand cleavage had an amplitude of 8.9 nM and data for 2^nd^-strand cleavage had a rate of 0.007 min^−1^.

Previous reports have shown that when 5’RNA and the 3’UTR are both present in a cleavage assay, 2^nd^ strand cleavage is inhibited ^6,7^. We examined the kinetics of cleavage when both 3’ and 5’RNA are present by using L-RNA which contains both 3’ and 5’RNA sequences on the same RNA molecule separated by a short spacer (**Figure 1B, Table 3**). As shown in **Figure 3C**, the amount of 1^st^ strand cleavage stimulated by L-RNA was comparable to that seen with 3’RNA or 5’RNA alone. However, the amplitude of 2^nd^ strand cleavage was lower with L-RNA than with 5’RNA (4 vs 14 nM, respectively), but occurred at a similar rate to that with 5’RNA alone (0.02 vs 0.05 min^−1^ for L-RNA vs 5’RNA, respectively). In a trapping experiment examining 1^st^- and 2^nd^-strand cleavage with L-RNA (**Figure 3D**), no additional 1^st^ strand cleavage occurred after addition of the trap (similar to the result with 5’RNA alone, **Figure 3B**), but unlike the data with 5’RNA no 2^nd^ strand cleavage occurred after the trap was added. These experiments confirm that 2^nd^ strand cleavage requires 5’RNA and can be inhibited by 3’RNA. The enzyme-DNA complex in the presence of 5’RNA is not kinetically stable before 1^st^ strand cleavage but is stable after 1^st^ strand cleavage. This stability after cleavage enables 2^nd^ strand cleavage to occur slowly. In the presence of both 3’RNA and 5’RNA the DNA dissociates even after 1^st^ strand cleavage resulting in little 2^nd^ strand cleavage (**Figure 3D**).

### TPRT kinetics following 1^st^-strand cleavage

After measuring the kinetics of 1^st^- and 2^nd^-strand cleavage, we next looked at the TPRT reaction where the 1^st^-strand cleavage product primes reverse transcription of the 3’RNA template. In an initial attempt to bypass the cleavage step and directly measure the rate of polymerization, the R2 enzyme was incubated with 3’RNA and a pre-cleaved 1^st^-strand substrate for 30 minutes to allow equilibration of the enzyme, RNA, and DNA. All four dNTPs were then added to initiate the reaction. The PAGE gel is shown in **Figure 4A** and the quantitative data are plotted in **Figure 4B**. Extension to the full length 278 nt product was visible after 0.75 minutes. Fitting the data by simulation estimated an average rate constant for polymerization of 6.8 nt s^−1^ (40 s to reach the half amplitude midpoint).

**Figure 4:**
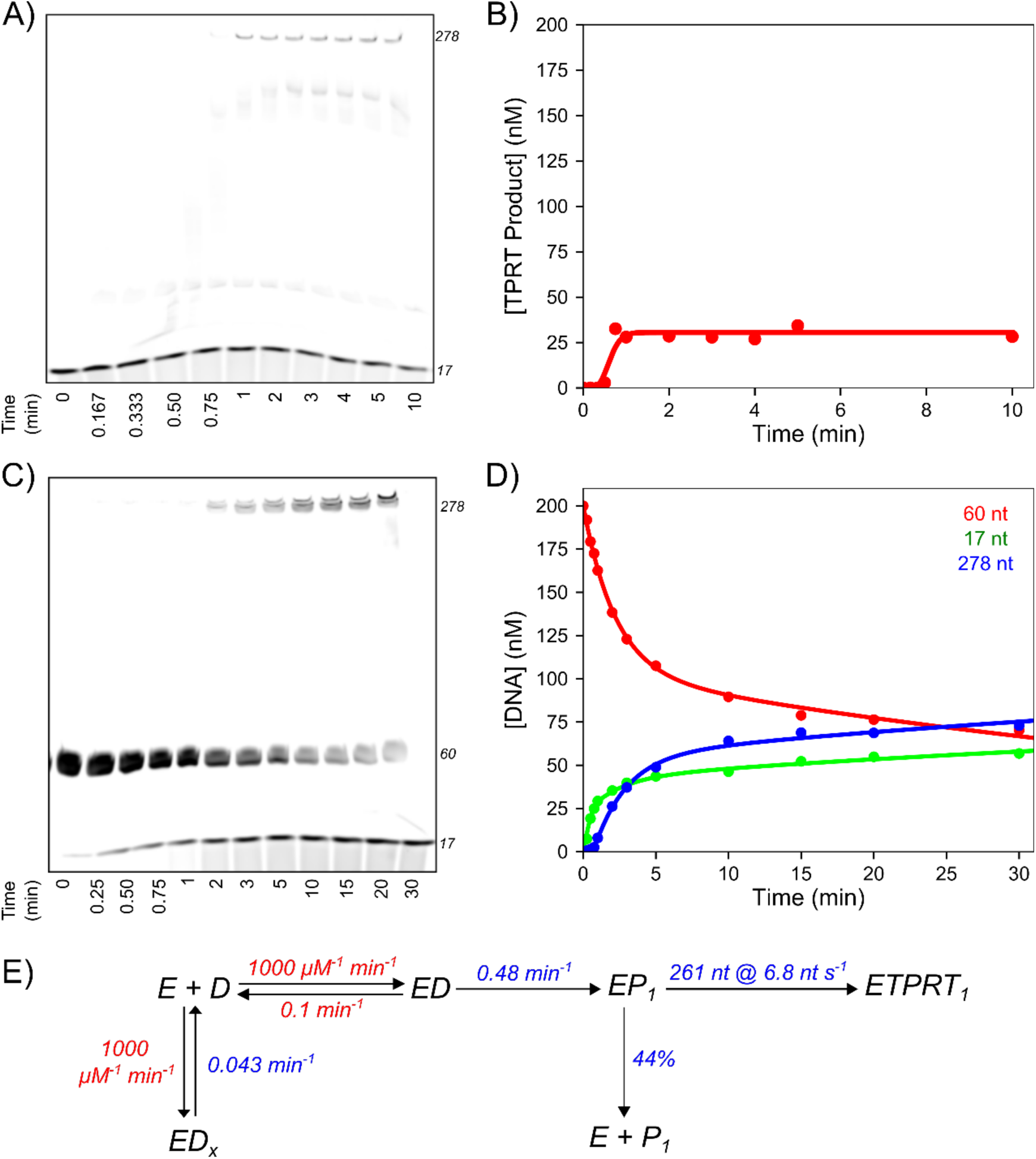
First strand cleavage and TPRT kinetics. <u>A) TPRT kinetics—PAGE.</u> A solution of 344 nM R2Bm, 1 µM 3’RNA, and 200 nM pre-cleaved DNA was mixed with 200 µM of each dNTP to start the reaction. <u>B) Quantification of TPRT kinetics and fit by simulation.</u> Data in A were quantified and fit by simulation in KinTek Explorer to define the kinetics of formation of the 278 nt product. <u>C) First strand cleavage and TPRT kinetics.</u> A solution of 344 nM R2Bm and 1 µM 3’RNA was mixed with 200 nM DNA and 200 µM of each dNTP to start the reaction. <u>D) Quantification and fit by simulation of 1^st^-strand cleavage</u> <u>and TPRT kinetics.</u> Data were fit by simulation in KinTek Explorer, where the red line represents the 60 nt starting material, the green curve represents the 17 nt cleaved product, and the blue represents the 277 nt TPRT product. <u>E) Kinetic scheme for cleavage and TPRT with 3’RNA.</u> Rate constants in red were locked at reasonable estimates that did not affect the shape of the curve over a wide range in the fitting. Rate constants in blue were derived by global data fitting. Confidence contours and best fit parameters are given in Figure S1 and Table S1, respectively.

We next measured the kinetics using a DNA that was not pre-cleaved, attempting to assess whether there is a kinetically significant step between cleavage and the start of polymerization, including the rate of transfer of the DNA from the endonuclease to the polymerase site of the protein. In this experiment, R2 enzyme and 3’RNA were pre-equilibrated. DNA and dNTPs were then added to initiate the reaction. **Figure 4C** shows the PAGE analysis demonstrating the cleavage of the 60 nt DNA to form the 17 nt primer, followed by TPRT primer extension to get the full length 278 nt product. The kinetics of the individual reaction steps are linked and difficult to resolve by traditional equation-based data fitting methods. Therefore, in **Figure 4D** we show analysis based on numerical integration of the rate equations and global data fitting^12,13^ to reveal the intrinsic rate constants summarized in **Figure 4E**. Confidence contour analysis established that each fitted parameter (shown in blue) was well constrained by the data (**Figure S1**). Note the lower amplitude of the TPRT product with the pre-cleaved DNA (**Figure 4B**) may reflect less efficient binding of the protein to an already cleaved DNA.

In the quantification, the cleaved 17 nt DNA product appeared first, then there was a short lag due to the sequential incorporation of 261 nt to reach the full-length 278 nt TPRT product. Fitting the data by simulation gave a rate of cleavage of 0.48 min^−1^ (0.008 s^−1^) followed by an average polymerization rate of 6.8 nt s^−1^, the same rate obtained in **Figure 4B**. These calculations do not include a step for transfer of the cleaved DNA primer from the endonuclease active site to the polymerase active site. This DNA transfer following cleavage therefore must be much faster than 0.008 s^−1^. Accordingly, DNA nicking (or a step before cleavage) is the rate-limiting step for 1^st^-strand TPRT synthesis. The DNA transfer reaction is moderately efficient as measured by the fraction of cleaved DNA that gets extended to the full-length product (56%). At the end of the transient phase (∼10 min) ∼60% of the DNA was cleaved, and ∼50% of that was extended to full length product. To account for the incomplete cleavage and extension, we propose a non-productive DNA binding state, which in the presence of dNTPs, slowly re-equilibrates to active state to give the slower linear increase in product after the burst phase observed in **Figure 4D**.

### Second-strand Synthesis and Full Reaction Kinetics

While 1^st^-strand cleavage, TPRT, and 2^nd^-strand cleavage were demonstrated *in vitro* over 30 years ago by Luan et al^2^, 2^nd^-strand synthesis in which the single strand DNA formed during TPRT is copied is postulated to be the final step to complete the formation of duplex DNA, but has never been seen *in vitro*^14,15^. To investigate whether our enzyme preparation could catalyze 2^nd^-strand synthesis we set up a reaction of R2 enzyme using CM-RNA (complete minimal RNA), which differs from L-RNA in that it also contains the 5’-UTR (**Figure 1B, Table 3**). We also added a 3’ phosphate blocking group to prevent nonspecific DNA synthesis from the free 3’ ends of the DNA substrate, since this has been previously observed with the R2 enzyme^16^. Different fluorescent labels were added to the 5’ends of both target DNA strands to allow quantification of the kinetics of cleavage and extension for both strands in a single reaction. We placed Cy3 on the “bottom” (anti-sense) strand that primes TPRT and FAM for the “top” (sense) strand that primes 2^nd^-strand synthesis (**Figure 1C**). Reaction products formed as a function of time were resolved by denaturing PAGE, the individual products quantified, and data fit by simulation in KinTek Explorer using the minimal model shown in **Figure 5E**. As shown in **Figure 5A** and **Figure 5B**, cleavage of the first strand (60 nt) went to ∼90% completion, forming the 17 nt product with a rate constant of 0.17 min^−1^ (0.003 s^−1^). Cleavage was followed by TPRT at an average rate of 9.5 nt/s for the addition of 1426 nt, giving a net half-life for full length product formation of 2.5 min. As discussed in the previous TPRT kinetics section, this polymerization rate may be a slight underestimate due to the length of time required for the DNA target to move from the endonuclease to the reverse transcriptase active site. However, because this DNA strand transfer reaction appears to be much faster than the observed cleavage and polymerization reactions, the data could be fit globally without including the strand transfer step.

**Figure 5:**
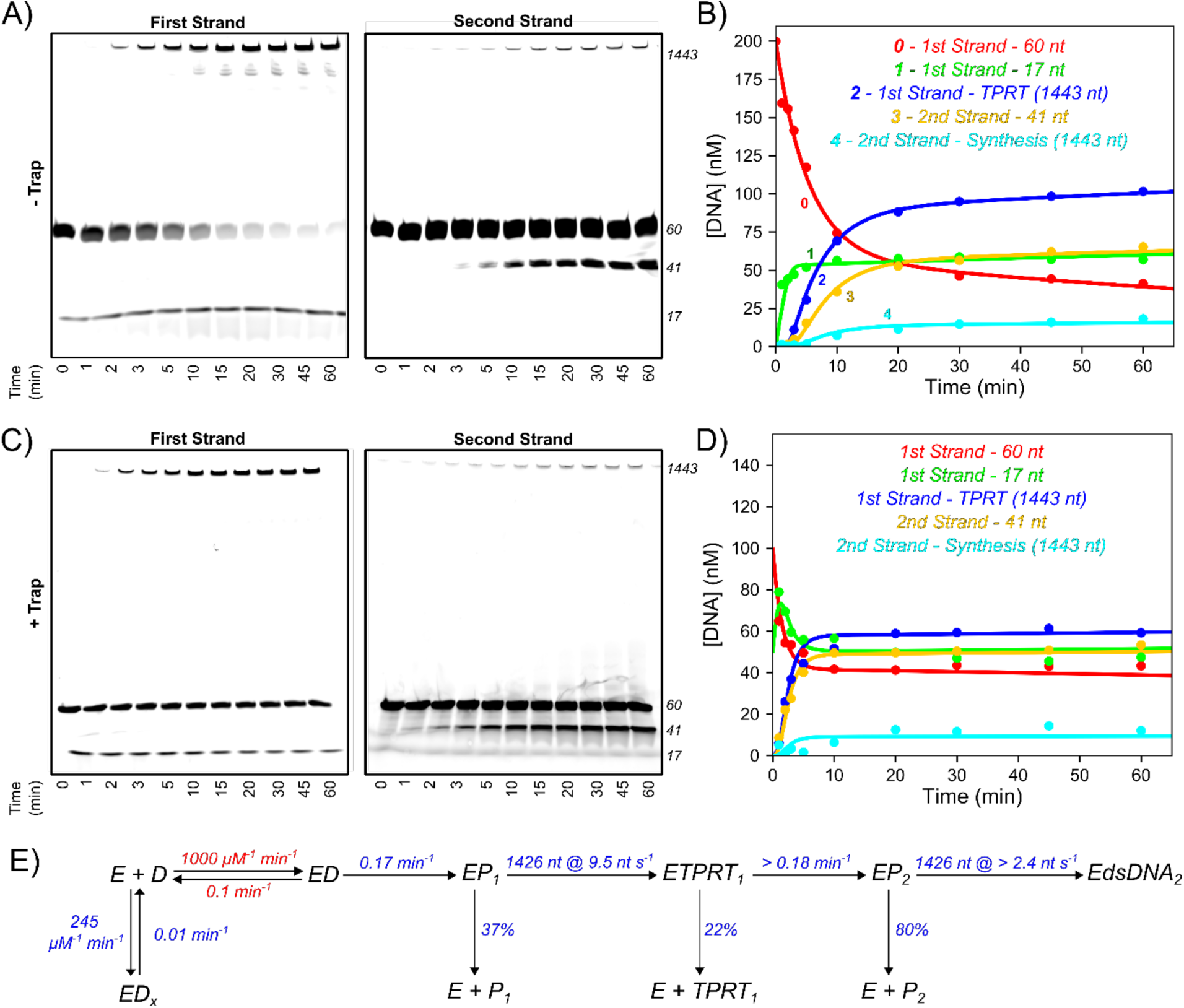
Reaction with Complete Minimal (CM) RNA. <u>A) Kinetics of full retrotransposition reaction by</u> <u>R2Bm.</u> A solution of 344 nM R2Bm and 1 µM CM-RNA was mixed with 200 nM DNA and 200 µM of each dNTP to start the reaction. PAGE for 1^st^ strand cleavage is shown on the left and for 2^nd^ strand cleavage is shown on the right. <u>B) Quantification and fit by simulation of full retrotransposition reaction.</u> Data were fit by simulation in KinTek Explorer to give the model shown in panel E. <u>C) Kinetics of full retrotransposition</u> <u>reaction by R2Bm, with DNA trap</u>. A solution of 387 nM R2Bm and 1 µM CM-RNA was mixed with 300 nM DNA and 200 µM dNTPs to start the reaction. After 1 minute, the reaction was diluted at a 1:1 ratio with unlabeled DNA to get a final concentration of 1 µM unlabeled DNA trap. The gel for the first strand is shown on the left and the gel for the second strand is shown on the right. <u>D) Quantification of full retrotransposition</u> <u>reaction with trap</u>. Data were fit by simulation in KinTek Explorer. <u>E) Reaction scheme for full</u> <u>retrotransposition reaction by R2Bm.</u> Rate constants in blue are from global fitting of data in A-B in KinTek Explorer. Rate constants in red were locked at reasonable estimates based on global data fitting. Confidence contours and best fit parameters are given in Figure S2 and Table S2, respectively. Confidence contours and best fit parameters for the reaction with unlabeled DNA trap are given in Figure S3 and Table S3, respectively.

Second-strand cleavage to form a 41 nt product was observed on a time scale mirroring the appearance of the TPRT product and was followed by rapid formation of the 2^nd^-strand synthesis product (**Figure 5A**). Quantitative analysis and global data fitting (**Figure 5B. 5E**) revealed that approximately 20% of the 2^nd^-strand cleaved DNA was extended to form the full-length 2^nd^-strand synthesis product. This contrasts with ∼60% completion of TPRT from the 1^st^ strand cleavage product. Although the efficiency of 2^nd^-strand synthesis is low, this represents the first direct observation of 2^nd^-strand synthesis and enables the first quantitative analysis to define the kinetics of the reaction. As with the data in **Figure 4**, confidence contour analysis shows that each fitted parameter (shown in blue) is well constrained by the data (**Figures S2** and **S3**). The DNA binding and dissociation steps (show in red) are not well defined and were locked at values sufficient to provide a minimal estimate.

Next, we performed a trapping experiment to quantify what fraction of the enzyme-bound cleaved DNA that partitioned forward to form the 2^nd^-strand synthesis product. R2 enzyme and CM-RNA were pre-incubated, then mixed with labelled DNA substrate and dNTPs to start the reaction (**Figures 5C-D**). After 1 minute, a large excess of unlabeled DNA was added to the reaction mixture to trap any free enzyme. The fraction of the DNA that was cleaved after adding the trap was reduced only slightly, indicating that the DNA does not readily dissociate from the enzyme prior to 1^st^-strand cleavage. In the presence of the trap, 60% of the 1^st^-strand cleaved DNA went on to form the TPRT product, followed by 2^nd^ strand cleavage and 2^nd^-strand synthesis. The observation of 2^nd^-strand synthesis in the presence of the trap suggests that (∼10%) of the initial protein complex stays bound to the RNA/DNA throughout the entire integration reaction. If a complete integration requires two subunits of the R2 protein, as has been proposed^6^, then both subunits appear able to remain bound to the target DNA. The rate of 1^st^ and 2^nd^ strand cleavage, but not the amount, increased by ∼3-fold in the presence of the DNA trap (**Table S2-S3**) but the rate of TPRT was unaffected by the trap. How and why these rates are affected by excess DNA is unknown but could be an interesting topic of investigation for future studies.

While the kinetics of appearance of the products demonstrated direct observation of 2^nd^-strand synthesis, we next aimed to verify that 2^nd^ strand synthesis had indeed been observed as opposed to formation of a nonspecific reaction product (**Figure 6, Figure S4**). A complete reaction was performed as in **Figure 5A-B**, the reaction was stopped, the mixture was treated with RNase, and then the DNA products were purified. PCR with one primer in the 28S region upstream of the insertion site and another primer in the 5’UTR gave rise to a strong band of the expected size (approximately 230 bp, within the resolution of the gel, **Figure 6B**). Sanger sequencing analysis showed good agreement of the expected PCR product in the sequence leading up to the 28S/5’UTR junction (**Figure 6C**). These data demonstrate accurate 2^nd^-strand synthesis to provide a cDNA copy of the 1400 nt RNA. Close analysis revealed multiple sequencing peaks starting at the 28S/5’UTR junction suggesting a mixed population of start sites for second strand synthesis or indels (**Figure 6C**).

**Figure 6:**
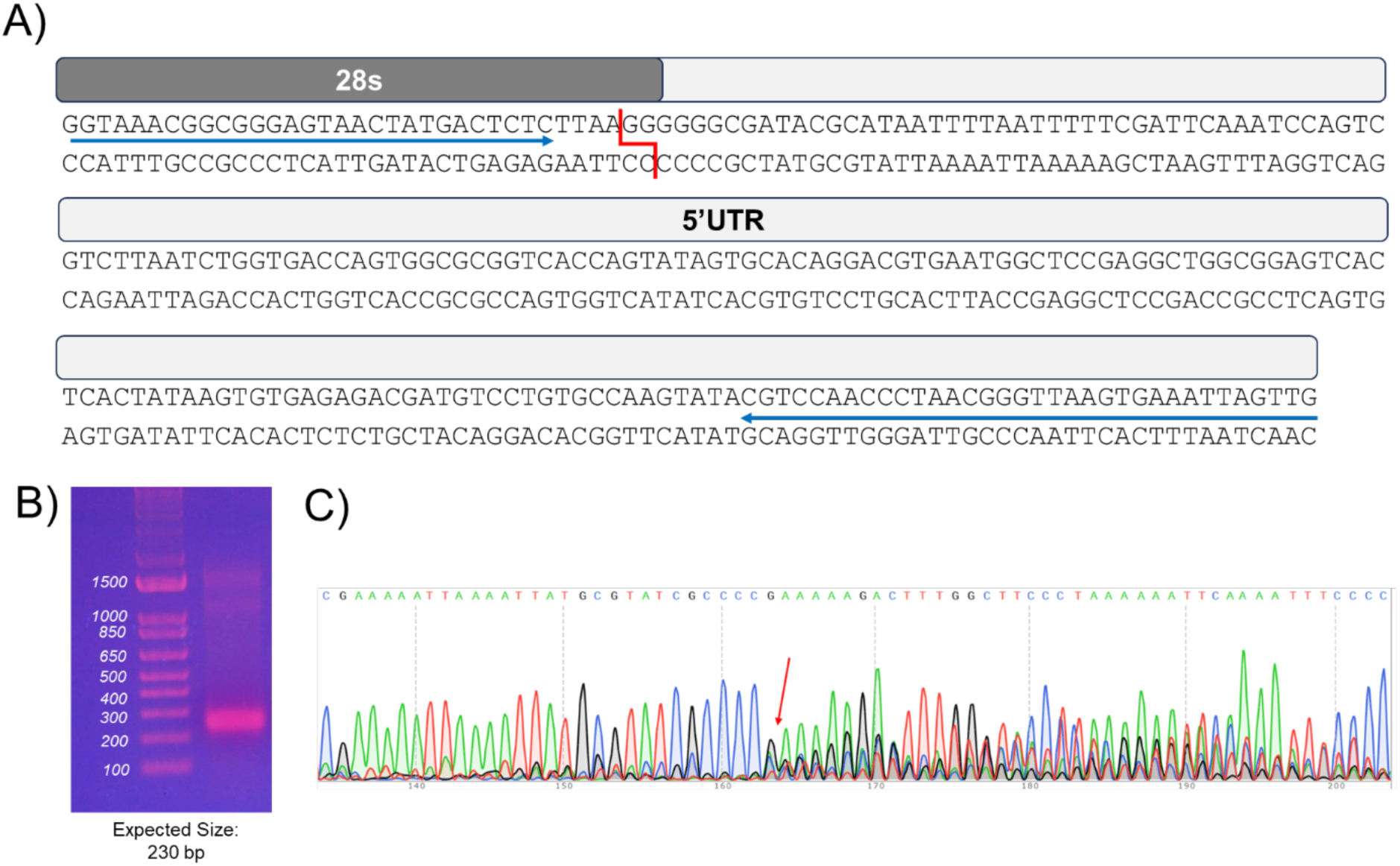
Analysis of 5’ Junction. <u>A) Diagram of 5’ junction PCR product</u>. Primer binding sites are given by the blue arrows. The cleavage sites are shown in red. The dark grey and light grey boxes show the 28s and 5’UTR portions of the PCR product, respectively. <u>B) PCR product using TPRT product as a template</u>. Expected size of the PCR product is approximately 230 bp. <u>C) Sanger sequencing analysis of 5’ junction</u>. The reverse primer was used for Sanger sequencing on the PCR product. Sequences up to the top line in (A) were omitted to better focus on the sequence at the junction. Bases are well resolved until the 28s junction, show with the red arrow. The overlapping peaks signify a mixed population of indels at this junction, as confirmed by Next Generation Sequencing amplicon variant analysis (see **Figure S4**).

Amplicon sequencing of the PCR product by next generation sequencing confirmed that the PCR product contained a variety of slightly different junction sequences (**Figure S4**). There were two locations where 2^nd^-strand synthesis appeared to originate: 52% of the junctions initiated near the 2^nd^-strand DNA cleavage site while 48% of the products initiated 2^nd^-strand synthesis at the 3’ end of the target DNA top strand, implying that the DNA polymerase initially copies the local DNA template before switching to the cDNA formed during 1^st^-strand TPRT. Junction sequences of each type are summarized in **Figure S4**. Of those originating near the cleavage site, some junctions had 1 – 3 additional residues (usually G or T residues), suggesting the enzyme can added a few non-templated bases before engaging the R2 cDNA sequences. Some junctions contained variable deletions of the R2 sequences, suggesting that reverse transcript had not gone to the end of the RNA template, or that the polymerase initiated second strand synthesis at an internal site on the cDNA. Such junctions would correspond to the 5’ truncations seen in many endogenous non-LTR retrotransposon insertions ^17^. Finally, some low abundance junctions were missing one to three bases upstream of the previously reported top strand cleavage site, suggesting that under the conditions of our assay, second strand cleavage may be somewhat variable. Surprisingly, slightly less than half of the junctions (∼48%) contained the downstream sequence of the 28S DNA substrate before R2 sequences were detected (**Figure S4**, bottom panel of junctions).

Because quality control analysis of our 3’ phosphate blocked DNA substrates showed almost complete 3’ phosphate modification (**Figure S5**), we can rule out nonspecific polymerization from the 3’ end of the DNA substrate^16^. Instead, the second strand appears to have been cleaved, then for some products the starting 28S bottom strand was copied before engaging the cDNA to polymerize the R2 sequence. For these sequences that contained downstream 28S sequence, the jump to the R2 cDNA sequences again resulted in 5’ truncated sequences. The variation seen in these 5’ junction sequences confirm that second strand synthesis was indeed detected in our *in vitro* assay and that the junctions generated had some of the characteristic seen in endogenous R2 insertion events as well as other non-LTR retrotransposons^17^. The use of R2 RNA containing 20 bases of the upstream 28S sequence did not change the kinetics or the amplitude of the complete integration reaction.

## Discussion

To survive as a parasitic gene in a host genome, mobile elements cannot generate excessive numbers of new insertions. As a result, their integration mechanism is probably not under selective pressure to be highly efficient. Non-LTR retrotransposons are a great example of this as their insertion seems quite inefficient, frequently generating variable 5’ and 3’ ends and extensive 5’ truncations. For a non-LTR retrotransposon to be engineered to create an efficient gene insertion tool, we need to begin with accurate characterization of the integration reaction with emphasis on the rate limiting steps as well as where the variability arises. The difficulties of working with purified non-LTR retrotransposon proteins is well supported by the limited number of publications on this topic, even though the first steps of the integration was documented nearly 30 years ago^2^. In general the *in vitro* activity of purified R2 proteins has been low and unstable^2,3,18^ Recently great success has been made in obtaining active enzymes *in vitro* and these studies, along with cryo-EM structural determinations, have greatly expanded our knowledge of R2 protein structure and activities ^7,8,10,14,15^. Success has also been obtained with new transformation assays using the R2 element found in a number of different species to further define the integration steps^7,15^. However, difficulties studying the protein *in vitro* remain and a wide range of activity levels have been reported with 1^st^-strand cleavage occurring on the order of minutes to hours^7,8,19^. Different tags, purification methods, reaction conditions, and R2 variants have been used and can partially explain these differences in activity. Presumably significant differences in the fraction of active enzyme in each preparation is also a contributing factor.

Here we present comprehensive analysis the kinetics of the individual steps of a complete *in vitro* insertion reaction using a relatively active enzyme preparation. To meet the standard for quantitative biochemistry in modern enzymology, we began by performing an active site titration. This type of experiment has been critical to examine DNA polymerase kinetics^20^, Cas9 kinetics,^11^ and many other systems where the active fraction is significantly less than 100%. Our assay relies on a fast rate of R2 catalysis relative to dissociation from the substrate and product DNA after initial binding so that the enzyme-DNA equilibrium is essentially fixed on the time scale of the activity measurement. Our data also show that the enzyme-DNA complex is kinetically stable and binding is tight (K_d_ < 4 nM). However, we found that only approximately 40% of our enzyme bound to the target DNA, and approximately 1/4 of that enzyme-DNA complex was in a nonproductive state. Such low activity may explain why a complete integration reaction was not detected in previous experiments even though a large excess of enzyme over DNA was used to saturate binding.

The rates of 1^st^- and 2^nd^-strand cleavage by the R2 enzyme are stimulated by different RNA molecules^6,7^. With just the 3’RNA, only the first strand is effectively cleaved, at a rate of approximately 0.6 min^−1^. With just 5’RNA, a segment of the 5’ end of the R2Bm ORF (**Figure 1B and Table 3**), both first and second strands of the DNA are cleaved with the first strand cleaved at an initial rate approximately 5 times faster than that of the second strand cleavage. When we tested the stability of the 2^nd^-strand cleavage product with a trapping experiment, we found that DNA was loosely bound before cleavage. This contrasts with the experiment with 3’ RNA in which the protein is stable before cleavage. Similarly, with L-RNA, containing the 5’RNA region through the 3’UTR, we found that most of the DNA dissociated from the enzyme before it was cleaved. Why the system evolved to have such stark differences in kinetics for first and 2^nd^ strand cleavage is likely due to the need for efficient 1^st^-strand cleavage and fast TPRT to prevent the formation of double stranded DNA breaks by premature 2^nd^ strand cleavage.

The TPRT reaction is quite complicated. R2 protein not only cleaves DNA but also uses the nicked DNA as a primer for TPRT. Because only 3’ RNA is reverse transcribed the enzyme must uses the 3’ RNA as a cofactor that binds to an allosteric enzyme site, and then this bound RNA must unfold to allow reverse transcription. Thus, it is important to understand the kinetics for the 1^st^-strand cleavage and the subsequent reverse transcription. We monitored the TPRT reaction using both pre-cleaved and uncleaved DNA substrates. We found no evidence for a kinetically significant step representing migration of the DNA from an endonuclease site to the polymerase site, indicating that the rate of transfer between the two sites is faster than DNA cleavage.

We also measured the rate of polymerization by using a pre-cleaved DNA substrate, preincubated with R2 and 3’RNA, and dNTPs were then added to start the reaction. An average rate of polymerization of 6.8 nt/s was observed, which is on par with HIV reverse transcriptase ^21^. Furthermore, TPRT involves polymerizing through large stretches of secondary structure in the 3’UTR and 5’RNA/5’UTR segments without leading to pausing and accumulation of significant intermediate states. Only minor pausing is seen towards the end of the RNA template leading to a small fraction of the paused intermediate prior to forming the full length TPRT product (**Figure 4A**). While we have not directly measured processivity, which is a function of the rate of polymerization divided by the rate of dissociation of the DNA from the enzyme, we see mostly complete extension to form the full-length cDNA with few intermediates that would accumulate due to dissociation of the DNA from the enzyme. A previous study also confirmed the high processivity of the enzyme and that it is not slowed by secondary structure^22^. Future studies could more accurately measure the processivity of this enzyme with longer RNA templates and trapping experiments.

To further understand the impact of RNA on the enzyme activity, we set out to test the enzyme with “complete minimal RNA” (CM-RNA), containing all structured RNA elements from the 5’UTR through the 3’UTR (**Figure 5A-5B**). First strand cleavage was efficient and was followed by formation of full length TPRT product at an average polymerization rate of around 9 nt/s. Second strand cleavage then followed at a rate of at least 0.18 min^−1^. After 2^nd^ strand cleavage, we observed rapid 2^nd^-strand DNA synthesis again leading to full length product. While we were not able to determine a polymerization rate directly due to the small amplitude of the 2^nd^-strand synthesis, our data fitting puts a lower limit of 2.4 nt s^−1^ on the rate of 2^nd^-strand synthesis. It should be noted that this rate involves displacement of the RNA still annealed to the cDNA. Approximately 80% of the DNA dissociated after the 2^nd^-strand cleavage and did not make it to the 2^nd^-strand synthesis, suggesting that this step significantly limits the efficiency of the overall insertion reaction. The initial 1^st^ strand DNA cleavage step is still more likely to represent the specificity-limiting step in the overall process since once the DNA is cleaved, the enzyme does not dissociate.

To test whether a complete insertion event occurs in a single DNA binding event, we performed a trapping experiment (**Figure 5C-5D**), as we performed for the other RNA substrates described earlier. In the trapping experiment, we observed that the rate of 1^st^-strand cleavage was around 3-fold faster than the reaction without the trap, so that the overall trap reaction was faster than the reaction without the trap. Furthermore, the amplitude of 2^nd^-strand synthesis was comparable to the amplitude observed without the trap. This indicates that R2 bound to DNA in the initial complex can perform 2^nd^-strand synthesis without dissociating from the DNA substrate. Although it has been proposed that two subunits of the R2 enzyme are required to complete this multistep reaction, at this time we have no data to evaluate this model since all our experiments were performed with a large molar excess of enzyme relative to DNA.

Previously, 2^nd^-strand synthesis was not detected *in vitro*. To confirm that we had indeed measured 2^nd^-strand synthesis and exclude the possibility of non-specific products, we performed an analytical PCR where one primer was located in the upstream 28S region of the DNA template and the other primer was located in the 5’UTR region of the cDNA (**Figure 6**). We observed a band of the expected size around 230 bp (**Figure 6B**). We then analyzed this PCR product by Sanger sequencing (**Figure 6C**) which verified that the product corresponded to the template RNA up until the 5’ junction. indicating that the cDNA was indeed used as the template for synthesis. Amplicon sequencing of the PCR product revealed a small fraction of the products with a precise 5’ junction (i.e. the 2^nd^ strand cleavage site priming synthesis from the very end of the R2 sequences). The remainder of the products were variable and contained many of the properties seen at the 5’ end of endogenous R2 inserts^17^. These included short deletions of the upstream 28S sequences, non-templated nucleotides, and small to large deletions of the R2 sequences. This variation is at least partly explained by the requirement that the enzyme must engage the cDNA template without the benefit of base pairing as the cDNA made by TPRT cannot anneal to the upstream DNA. We have referred to this ability of the R2 protein to switch to another template without complementarity as “template jumping”.

An unexpected finding from the analysis of the 5’ products was that a template jump to the cDNA was preceded in nearly half the cases by the enzyme jumping to the lower 28S strand downstream of the cleavage site before engaging the cDNA. These “two-jump” junctions may be explained by the ease with which the downstream DNA, only 19 bp in length, can be denatured after second strand cleavage. Further experiments should help to resolve these critical steps and the most accurate templates to be used in a complete integration reaction.

While here we report rates of the reaction, other studies were recently published with some rudimentary kinetic analyses of 1^st^ and 2^nd^ strand cleavage. One paper assigned no rates to their cleavage reaction, but showed cleavage occurring after a 30 minute incubation^88^. Another paper did provide some rate estimates^7^ which are more than an order of magnitude slower than what we report here. None of the previous work observed 2^nd^ strand synthesis. It is notable that we are using a different reaction buffer, described by Wilkinson et al^8^, that despite being high in salt (400 mM potassium acetate), provided us the best activity when screened against other reaction buffers from the literature^3,7^. Of note, we also did not cleave the MBP-SUMO tag from the enzyme before our kinetics assays due to the instability of the protein after tag cleavage. In future work, it is worth exploring ways to stabilize the protein without tags and re-determine its kinetics.

The major achievement of this work is summarized in **Figure 5B** and **Figure 5E**, where global kinetic analysis of a single turnover of the enzyme cycle afforded estimates of intrinsic rate constants for each step in the sequential pathway. Because the steps are kinetically linked, conventional fitting of data using exponential functions fails to provide interpretable results. In particular, the kinetics of formation of reaction products 2, 3, and 4 (**Figure 5B**) show long lags due to the processive DNA polymerization that cannot be fit accurately using an equation. Our analysis shows that reverse transcription of the RNA (TPRT) is fast and processive with no significant intermediates, incorporating 1426 nucleotides at an average rate of 9.5 nt/s. For processive polymerization, the time to reach the midpoint of the curve can be estimated as the number of nucleotides (1426) divided by the rate constant per nucleotide (9.5 nt/s). However, this approximation does not account for the time required to complete 1^st^-strand cleavage which adds to the observed lag time. All of these features are rigorously modeled without approximations based on numerical integration of the rate equations to yield a single unifying model ^12,13,23^. For 2^nd^-strand synthesis the amplitude is small, and the reaction appears coincident with 2^nd^-strand cleavage, so our analysis can only provide a lower limit on the average rate constant for polymerization >2.4 nt/s. Nonetheless, the data show that 2^nd^-strand cleavage, which provides the primer needed for 2^nd^-strand polymerization, is rate-limiting.

Our model accounts for all the data (**Figure 5B**) and defines the sequential four step reaction: 1^st^-strand cleavage, TPRT, 2^nd^-strand cleavage and then 2^nd^-strand synthesis to produce duplex DNA. Using the CM-RNA, 2^nd^-strand cleavage and subsequent synthesis do not occur until after TPRT is completed. Although we could envision a model where two enzymes are bound in opposite orientations on the DNA to afford cleavage of the two strands^24^, we have no evidence to distinguish whether one or two enzymes are employed to complete the full reaction sequence because essentially all of our experiments were performed with enzyme in excess. However, the sequential reactions outlined in **Figure 5D** require coordination between the endonuclease and polymerase sites which may be better accomplished with a single enzyme, while a dimer would require allosteric coordination between the two subunits.

While we have observed the long-sought 2^nd^ strand synthesis in our experiments, many open questions remain about the R2Bm retrotransposition reaction. Recent structures provided images to describe 1^st^ strand cleavage and TPRT, but attempts to obtain high resolution structures of the 2^nd^ strand cleavage complex have failed. The transient nature of the interaction may preclude capturing this complex with R2Bm. While structural studies provide elegant snapshots of enzymes, they do not provide information on the underlying dynamics of enzyme-substrate interactions. Understanding the kinetics underlying an enzyme catalyzed reaction can greatly facilitate future structure determination by elucidating reaction conditions that can trap a particular state. Overall, the experiments described here lay a framework for future work on R2Bm and other R2 elements that will help to quantify enzyme activity assays, including 2^nd^ strand synthesis. Further understanding of the biochemistry of 2^nd^ strand synthesis reactions will accelerate our ability to engineer this limiting step for more efficient gene insertion by R2 elements and capture more structural states during second strand synthesis.

Our analysis reveals inefficiencies in the initiation of 2nd-strand synthesis in terms of fractional conversion of the TPRT product to duplex cDNA and in accurate selection of the cDNA at the start site for replication. Both aspects should be targeted in efforts to improve the overall efficiency of RNA replication since little is known about how the primer formed during 2nd-strand cleavage gets transferred to the polymerase site, or how the TPRT product is aligned in the template position to give accurate replication to form duplex cDNA. Despite the observed limitations in efficiency, it is remarkable that we have been able to observe a physiologically significant rate for accurate replication of a 1400 nt RNA to form duplex cDNA in approximately 15 minutes. There is no reason to believe this is an upper limit to insert size. Our data quantify the fundamental reactions catalyzed by an R2 non-LTR transposon and provide a path forward to engineer a more efficient enzyme with the potential to insert large genes in vivo.

## Materials and Methods

### Expression and Purification of R2Bm

R2Bm was expressed and purified using a previously described protocol^88^. R2Bm Δ1-110 was expressed and purified with a N-terminal 14x-His-MBP-SUMO tag and a C-terminal twinstrep tag (**Figure S6**). The plasmid pMW41 was ordered from Addgene (Plasmid #209,290, Watertown, MA), extracted, transformed into electrocompetent BL21(DE3) *E. coli*, plated on LB/agar/ampicillin (50 µg/ml) plates, and incubated overnight at 37°C. The following day a single colony was inoculated into a 100 ml starter culture of Terrific Broth (TB) (2.4% (w/v) yeast extract, 2% (w/v) tryptone, 0.4% (v/v) glycerol, 0.017 M KH_2_PO_4_, 0.072 M K_2_HPO_4_) + 50 µg/ml ampicillin. and incubated overnight with shaking at 32°C, 250 RPM. The following morning, a 1 L culture of TB Autoinduction Media (TB + 50 mM ammonium chloride, 5 mM sodium sulfate, 0.2% (w/v) α-lactose monohydrate, 0.05% (w/v) glucose, 2 mM magnesium chloride) + 50 µg/ml ampicillin. was inoculated to an OD_600_ of 0.1 and incubated with shaking at 37°C, 250 RPM until the OD_600_ reached 0.8. The culture was then moved to an incubator set to 22°C and grown for 16 – 20 hours with shaking at 250 RPM. The cells were then harvested by centrifugation at 6,000 x g for 20 minutes at 4°C. All subsequent purification steps were performed at 4°C or on ice. The cell pellet (∼15g) was resuspended in 100 ml Lysis Buffer (50 mM Tris-HCl pH 7.5, 1 M NaCl, 10% glycerol, 5 mM 2-mercaptoethanol). One EDTA Free protease inhibitor tablet (Roche) was added, and the cells were then lysed using a LM-20 microfluidizer (Microfluidics) with two passes through the instrument at 18,000 psi with ice cooling the interaction chamber and coils. The lysate was then clarified by centrifugation at 60,000 x g for 30 minutes at 4°C and filtered through a 0.2 µm PES syringe filter (Whatman). The clarified lysate was loaded onto a 5 ml StrepTactin SuperFlow Plus (Qiagen) gravity column equilibrated in Lysis Buffer. The column was washed with Lysis Buffer, then Wash Buffer (20 mM HEPES-KOH pH 7.9, 500 mM KCl, 10% glycerol, 5 mM 2-mercaptoethanol), and finally eluted with elution buffer (wash buffer + 5 mM desthiobiotin). Fractions were analyzed by SDS-PAGE and fractions containing R2Bm were pooled. Pooled fractions were concentrated to an A_280_ of 1.7 with a 50 kDa MWCO centrifugal concentrator (Amicon). The concentration of purified protein was determined by absorbance at 280 nm using the extinction coefficient 208,670 M^−1^ cm^−1^, calculated based on the amino acid sequence of the protein. Protein was flash frozen in small aliquots in liquid nitrogen and stored at −80°C.

### DNA Substrates

DNA oligonucleotides used in this study were purchased from Integrated DNA Technologies (IDT, Coralville, IA) with HPLC purification and resuspended to 100 µM in IDTE buffer (10 mM Tris-HCl pH 8, 0.1 mM EDTA). Oligos were annealed by mixing at a 1:1 molar ratio in DNA annealing buffer (10 mM Tris-HCl pH 7.5, 50 mM NaCl, 1 mM EDTA), heating to 95°C, then cooling slowly to room temperature over approximately 1 hour. Oligonucleotides were stored at −20°C. Sequences and extinction coefficients of oligos used in this study are shown in **Table 1**.

**Table 1:**
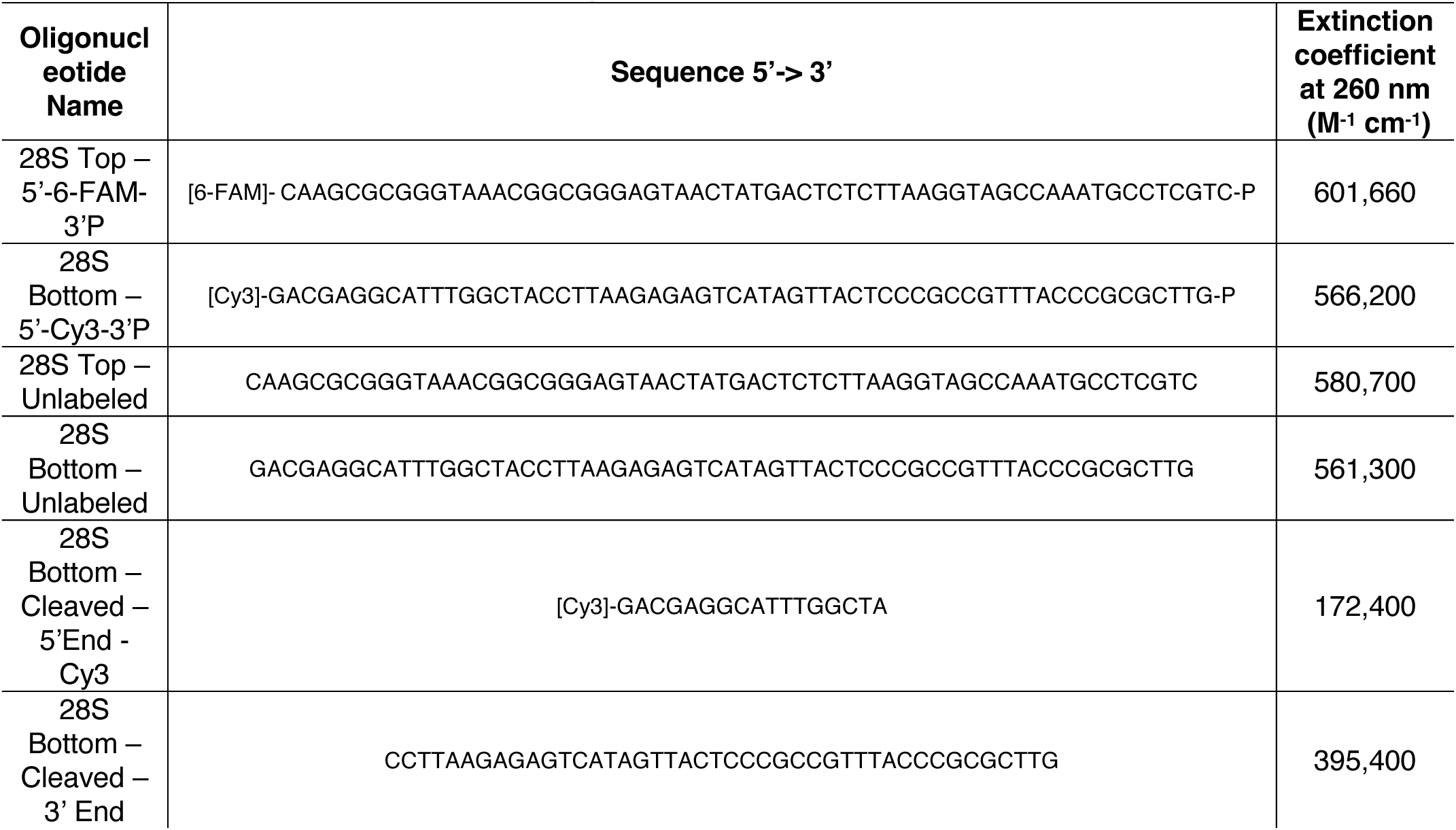
Oligonucleotides used in this study.

### RNA Preparation

A gene block to be used as a PCR template for in vitro transcription was synthesized by IDT containing the 5’UTR, a partial ORF containing the 5’RNA portion, and the 5’UTR. The sequence of the synthesized DNA is given in the supplemental information. mRNAs were prepared by in vitro transcription. Primers used to amplify regions of the gene block as templates for in vitro transcription are given in **Table 2**. PCR was performed with Q5 polymerase (NEB) with 5% DMSO as an additive then amplicons were purified with a PCR cleanup kit (Qiagen). In vitro transcription reactions were set up in a volume of 320 µl with 8 µg PCR product template DNA, 5 mM DTT, 2 mM NTPs, and 1600 Units of T7 RNA polymerase in T7 RNA Polymerase reaction buffer (NEB). Reactions were incubated at 37°C for 3 hours then digested with 64 units of RNase Free DNaseI (NEB) at 37°C for 15 minutes. EDTA was then added to 5 mM and the reaction was heat inactivated at 85°C for 10 minutes. RNA was then ethanol precipitated and dried to completion in a SpeedVac. RNA was resuspended in RNA annealing buffer (10 mM Tris-HCl pH 7, 50 mM NaCl, 0.1 mM EDTA), the concentrations were determined by absorbance at 260 nm using the extinction coefficients given in **Table 3**, and then the RNA was stored at −20°C.

**Table 2:**
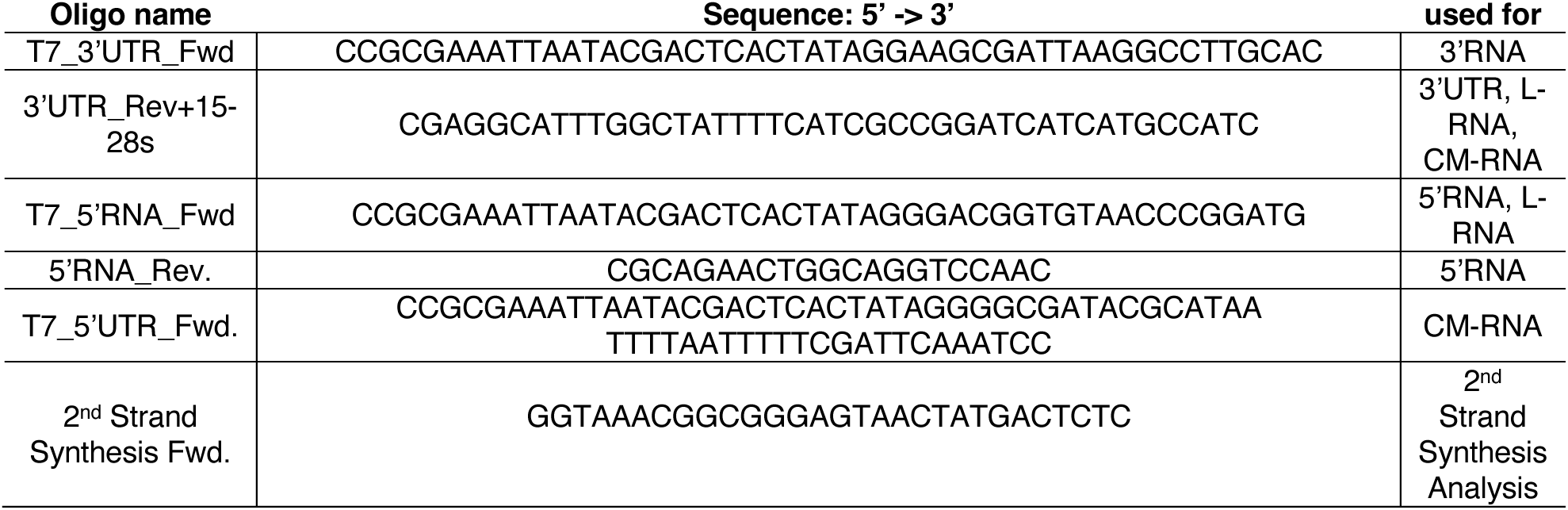
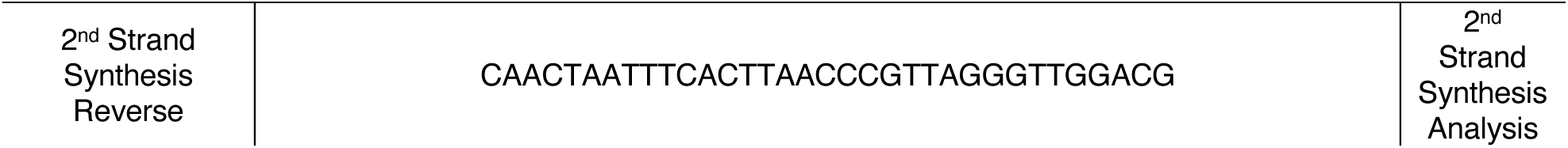
Primers used in this study.

**Table 3:**
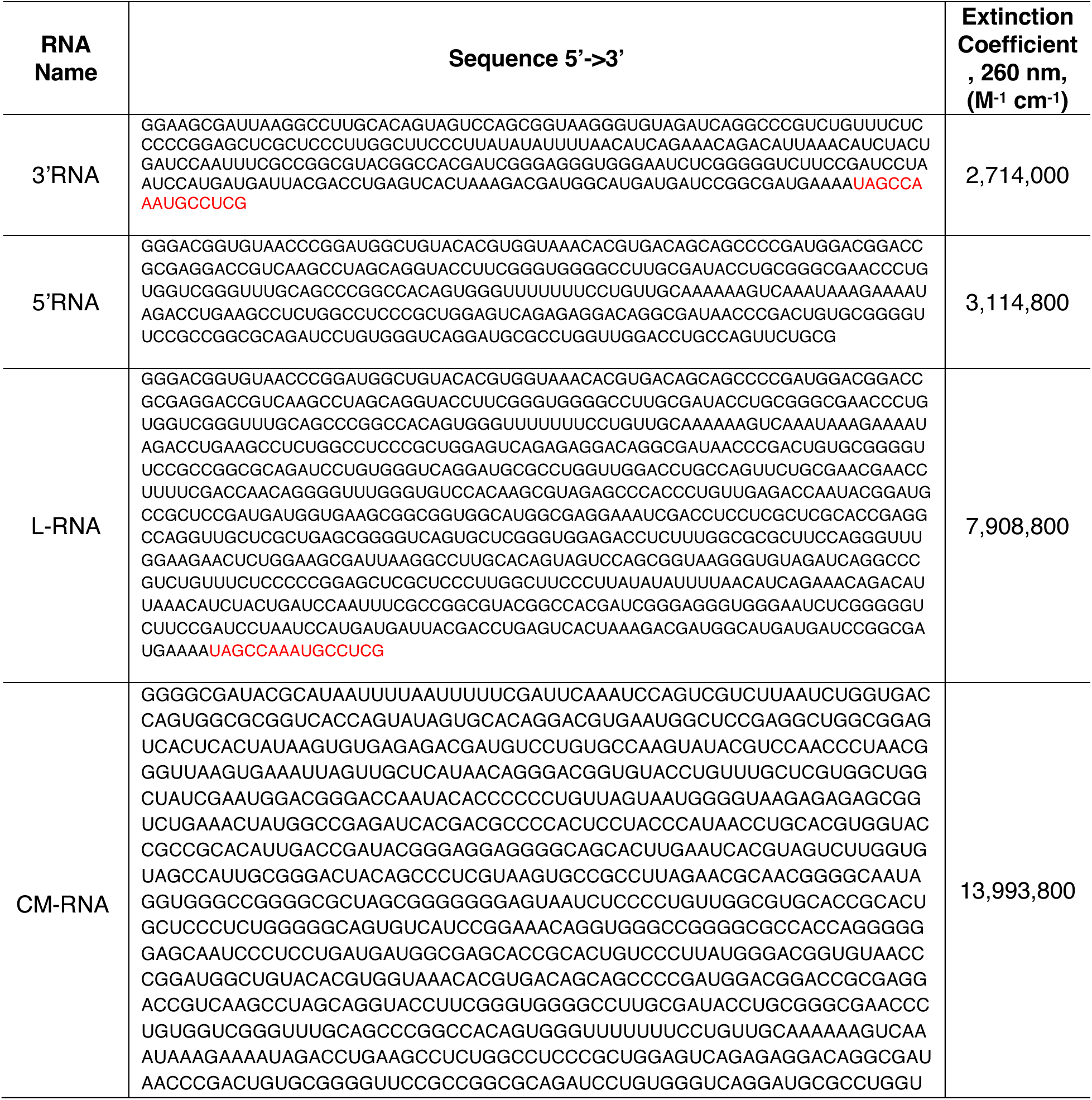
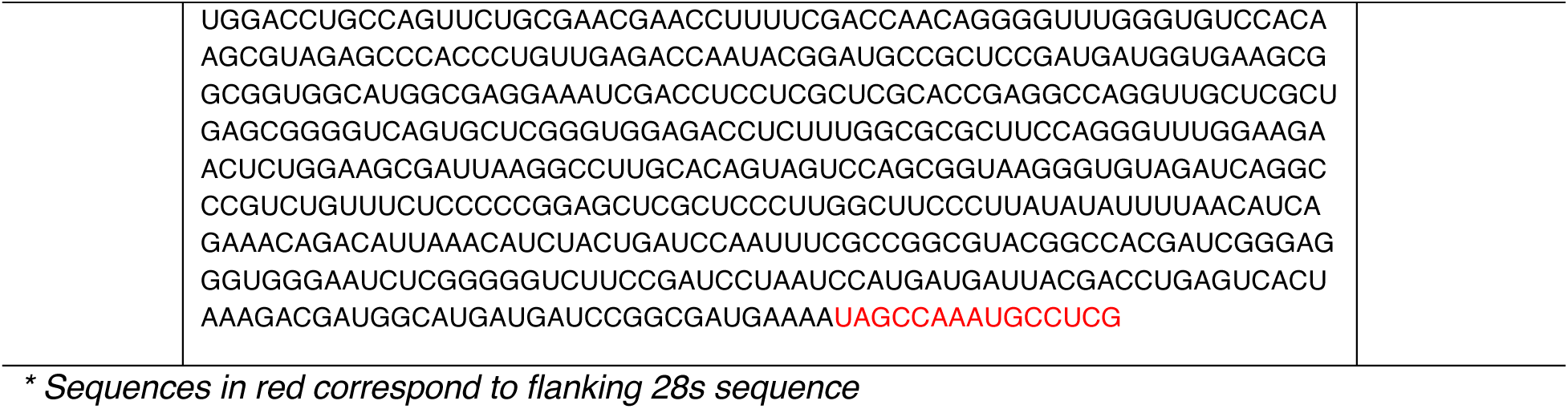
RNA used in this study.

### Kinetics Experiments

Unless otherwise noted, all experiments were performed in R2Bm Reaction Buffer + Mg^2+^ (20 mM HEPES-KOH pH 7.9, 400 mM Potassium Acetate, 5 mM Magnesium Acetate, 1 mM DTT). Reactions were set up with components at 2x their final concentration then mixed at a 1:1 ratio with substrate to start the reaction. For all reactions, RNA and protein were pre-incubated on ice for 30 minutes before performing cleavage/polymerization reactions. Concentrations given in the text are final concentrations after mixing, unless otherwise noted. Cleavage reactions were analyzed by capillary electrophoresis on an Applied Biosystems 3130xl instrument equipped with a 36 cm capillary array and nanoPOP-6 polymer (Molecular Cloning Laboratories, San Francisco CA) as previously described^252523^. Before analysis, 1 µl of quenched reaction time point was diluted into 10 µl of HiDi Formamide (ThermoFisher) and samples were injected at 3.6 – 5 kV for 10 – 60 seconds, depending on the concentration of labeled oligos in the reaction. Peaks were quantified with GeneMapper Software version 5. Representative electropherograms are given in **Figure S7** and **Figure S8**. Reactions analyzing TPRT or second-strand synthesis were analyzed by denaturing PAGE. Gels contained 12% acrylamide (19:1 acrylamide : bisacrylamide), 7 M Urea, and 1x TBE buffer (90 mM Tris-Borate, 2 mM EDTA) and were run in 1x TBE pre-heated to 55°C before loading samples and running the gel at 400V for approximately 20 minutes. Quenched samples were mixed at a 1:1 ratio with 2x TBE/Urea sample buffer (2x TBE, 7 M Urea, 12% Ficoll Type 400, bromophenol blue) and heated to 95°C for 5 minutes before loading on a gel. Gels were scanned on a Typhoon 9500 scanner (Cytiva). All reactions were performed at least twice on separate days to ensure reproducibility and representative data sets are shown in the paper. For all figures, except as noted in Figure 2, concentrations of enzyme given correspond to active R2Bm. Images were quantified with ImageJ software^2626^.

### Kinetics Data Analysis

Kinetic data were initially fit to equations using the *afit* function in KinTek Explorer. The equation for a single exponential function used was:

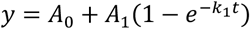

Where A_0_ is the y-intercept, A_1_ is the amplitude, *k_1_* is the decay rate and t is time. The equation for a single exponential burst equation used was:

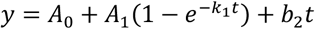

Where A_0_ is the y-intercept, A_1_ is the amplitude of the exponential phase, *k_1_* is the exponential decay rate, t is time, and b_2_ is the rate of the linear phase.

The equation for a quadratic function used was:

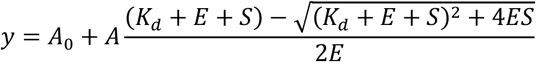

Where A_0_ is the y-intercept, A is the amplitude, *K_d_* is the dissociation constant, E is the enzyme concentration, and S is the substrate (DNA) concentration.

### Analysis of Kinetic Data in KinTek Explorer

Kinetic data were analyzed with KinTek Explorer v11.0.1 (KinTek Corporation, Austin TX)^23^. First a model was entered then each experiment was modeled by entering the starting concentrations of reactants used in the experiments as well as defining an observable output. In the models below, the equal sign defined the reversible reaction. Irreversible reactions were set by locking the reverse rate to 0 and these are indicated by the → symbol below. Parameters that were not well defined by the data were locked in the fitting and are listed in **Table S1** – **Table S3** and are shown in red in the reaction schemes in **Figure 4** and **Figure 5**. Values for the locked rate constants were obtained as the minimal value above which there are no changes in the observed reaction kinetics.

For the 3’RNA data in **Figure 4**, the model below was used. All rate constants in step 4 were linked and having multiple polymerization steps in sequence was necessary to account for the lag in TPRT product formation. The rate constant obtained for step 4 was divided by 10, then divided by 60 to give the rate constant in s^−1^, and multiplied by the number of nucleotides added to get the rate in nt s^−1^ reported in **Figure 4**.

1. E+D = ED
2. E+D = EDx
3. ED → EP1
4. EP1 → FP1 → FP2 → FP3 → FP4 → FP5 → FP6 → FP7 → FP8 → FP9 → ETPRT1
5. EP1 → E + P1

For the CM RNA data in **Figure 5**, the model below was used. All rate constants in step 4 were linked and having multiple polymerization steps in sequence was again necessary to account for the lag in TPRT product formation. The rate constant obtained for step 4 was divided by 10, then divided by 60 to give the rate constant in s^−1^, and multiplied by the number of nucleotides added to get the rate in nt s^−1^ reported in **Figure 5**. Binding and dissociation rates for the unlabeled DNA trap in step 9 were locked at 1000 µM^−1^min^−1^ and 0.1 min^−1^, respectively.

1. E+D = ED
2. E+D = EDx
3. ED → EP1
4. EP1 → FP1 → FP2 → FP3 → FP4 → FP5 → FP6 → FP7 → FP8 → FP9 → ETPRT1
5. ETPRT1 → EP2
6. EP2 → EdsDNA2
7. EP1 → E + P1
8. EP2 → E + P2
9. E+Trap = E.Trap

Confidence contours were calculated in KinTek Explorer using the FitSpace function of the software ^23^. For data in both **Figure 4** and **Figure 5**, the χ^2^ threshold corresponding to the 95% confidence interval (calculated based on the number of parameters and the number of data points in the fitting) was 0.8. Confidence contours are given in **Figure S1, Figure S2**, and **Figure S3**.

Calculations of Flux in comparing fraction of DNA that proceeded forward versus dissociated after 1^st^ and 2^nd^-strand cleavage were performed using a dynamic partial derivative analysis during data fitting based on numerical integration of the rate equations using KinTek Explorer software. Using the steps described above for the reaction with 3’RNA, looking at the partitioning of the EP1 species between dissociation and TPRT, we calculate the flux of enzyme using the syntax EP1::4 (flux of EP1 through step 4) for the flux through TPRT and EP1::5 for the flux through dissociation after 1^st^-strand cleavage. The percentage of enzyme that dissociates after 1^st^-strand cleavage is then 100*(EP1::5/(EP1::5 + EP1::4)).

### PCR Analysis of First and Second Strand Synthesis Reaction Product

A 100 µl reaction was assembled by mixing 0.344 µM R2Bm, 0.2 µM 28s DNA, 1 µM CM-RNA+3’28s, and 200 µM each dNTP and incubating at 37°C for 90 minutes. 2 µl of 20 mg/ml RNaseA (NEB) was added to the reaction and the reaction was incubated at room temperature for 5 minutes before quenching the reaction with 50 mM EDTA. The DNA reaction product was then purified using a Qiagen PCR cleanup kit. Purified DNA was used as a template in PCR with Q5 polymerase using the forward and reverse 2^nd^-strand synthesis analysis primers listed in **Table 2**. An aliquot of the PCR reaction product was analyzed on a 1.2% agarose gel (**Figure 6B**) and showed a product of the expected size (approximately 230 bp). The rest of the PCR product was purified with a Qiagen PCR cleanup kit. The purified PCR product was then used as a template with the 2^nd^-strand synthesis reverse primer for Sanger sequencing analysis (Eton Bioscience Inc., San Diego CA) (**Figure 6C**). Amplicon sequencing of the PCR product was performed by Quintara Biosciences (Cambridge, MA) and variant analysis was performed (**Figure S4**). Plasmid maps and sanger sequencing images were created with SnapGene® software (from Dotmatics; available at snapgene.com).

## Author Contributions

Conceptualization: T.L.D, K.A.J, J.Z. Investigation and data analysis: T.L.D. Writing initial draft: T.L.D. All authors reviewed, edited, and approved the manuscript. Supervision: K.A.J. and J.Z.

## Materials and Correspondence

Correspondence and material requests should be addressed to Kenneth Johnson and Tyler Dangerfield.

