## Supplemental Information for "Single-turnover kinetic analysis of non-LTR retrotransposition defines the mechanism and rate constants governing each step"

Sequence of R2Bm g block used for in vitro transcription

ggggcgatacgcataattttaatttttcgattcaaatccagtcgtcttaatctggtgaccagtggcgcggtcaccagtatagtgcacaggacgtgaatgg
ctccgaggctggcggagtcactcactataagtgtgagagacgatgtcctgtgccaagtatacgtccaaccctaacgggttaagtgaaattagttgctc
ataacagggacggtgtacctgtttgctcgtggctggctatcgaatggacgggaccaatacaccccccctgttagtaatggggtaagagagagcggtct
gaaactatggccgagatcacgacgccccactcctacccataacctgcacgtggtaccgccgcacattgaccgatacgggaggaggggcagcac
ttgaatcacgtagtcttggtgtagccattgcgggactacagccctcgtaagtgccgccttagaacgcaacggggcaataggtgggccggggcgcta
gcgggggggagtaatctcccctgttggcgtgcaccgcactgctccctctgggggcagtgtcatccggaaacaggtgggccggggcgccaccagg
ggggagcaatccctcctgatgatggcgagcaccgcactgtcccttatgggacggtgtaacccggatggctgtacacgtggtaaacacgtgacagc
agccccgatggacggaccgcgaggaccgtcaagcctagcaggtaccttcgggtggggccttgcgatacctgcgggcgaaccctgtggtcgggttt
gcagcccggccacagtgggttttttttcctgttgcaaaaaagtcaaataaagaaaatagacctgaagcctctggcctcccgctggagtcagagagga
caggcgataacccgactgtgcggggttccgccggcgcagatcctgtgggtcaggatgcgcctggttggacctgccagttctgcgaacgaacctttttc
gaccaacaggggtttgggtgtccacaagcgtagagcccaccctgttgagaccaatacggatgccgctccgatgatggtgaagcggcggtggcatg
gcgaggaaatcgacctcctcgctcgcaccgaggccaggttgctcgctgagcggggtcagtgctcgggtggagacctctttggcgcgcttccagggtt
tggaagaactctggaagcgattaaggccttgcacagtagtccagcggtaagggtgtagatcaggcccgtctgtttctcccccggagctcgctcccttg
gcttcccttatatattttaacatcagaaacagacattaaacatctactgatccaatttcgccggcgtacggccacgatcgggagggtgggaatctcggg
ggtcttccgatcctaatccatgatgattacgacctgagtcactaaagacgatggcatgatgatccggcgatgaaaa

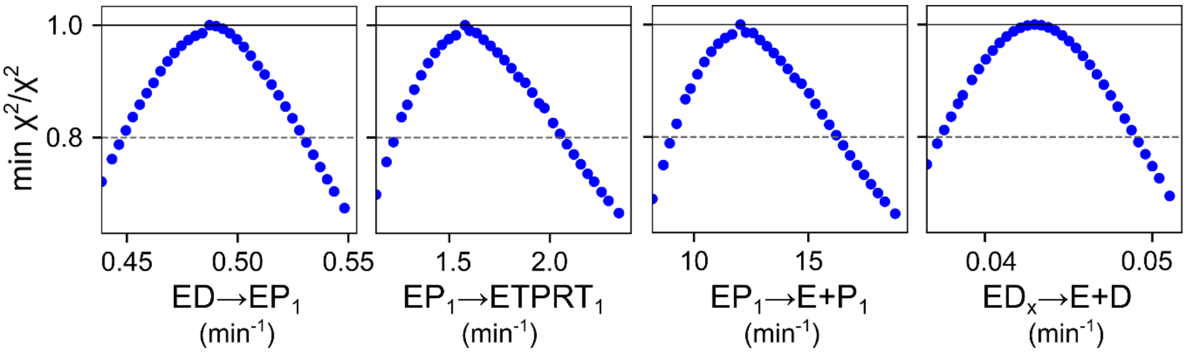


**Figure S1: Confidence Contours for First Strand Cleavage and TPRT Kinetics, Related to Figure 4**. Confidence contours were generated in KinTek Explorer by fitting to the scheme in Figure 4 (described in the methods section). The dashed line represents the $chi^2$ threshold (0.8) corresponding to the 95% confidence interval for estimated parameters. Best fit parameters and confidence intervals are given in Table S1.

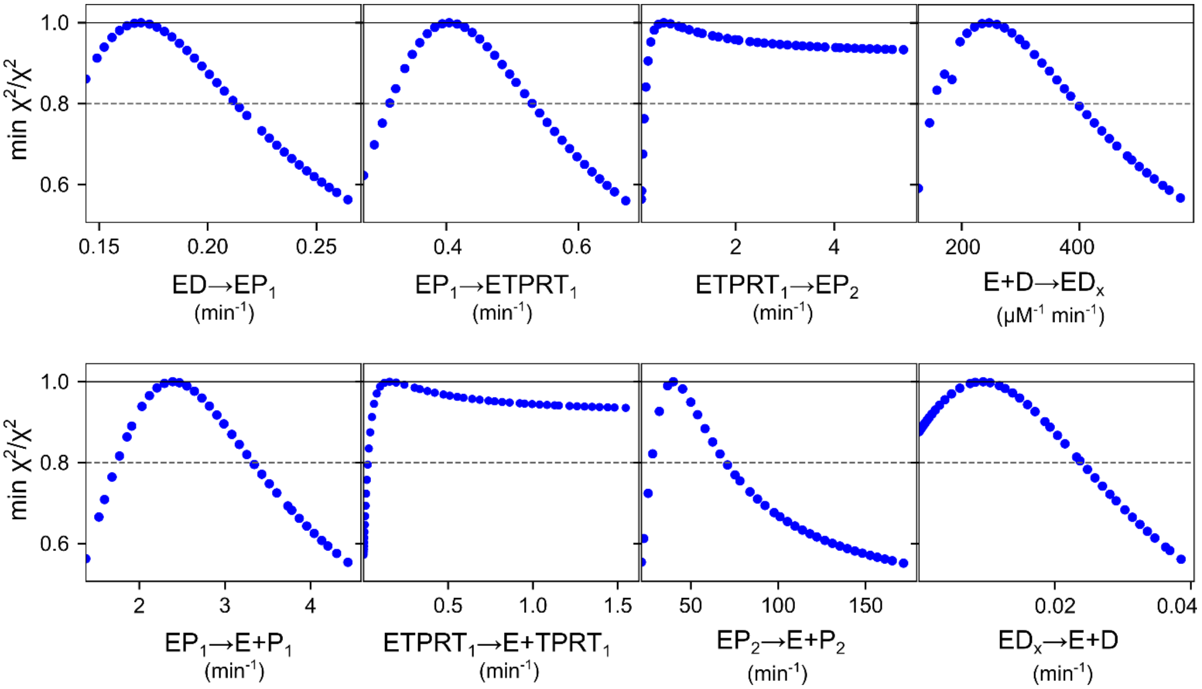


**<u>Figure S2: Confidence Contours for Kinetics of Reaction with Complete Minimal (CM) RNA.</u>** Confidence contours were generated in KinTek Explorer by fitting to the scheme in Figure 5 (described in the methods section). The dashed line represents the chi$^2$ threshold (0.8) corresponding to the 95% confidence interval for estimated parameters. Note that only lower limits are obtained for the $ETPRT_1$ ->$EP_2$ and $ETPRT_1$ -> E + $TPRT_1$ steps individually as the partitioning between dissociation and second strand cleavage is defined by the data rather than either rate constant individually. Best fit parameters and confidence intervals are given in Table S2.

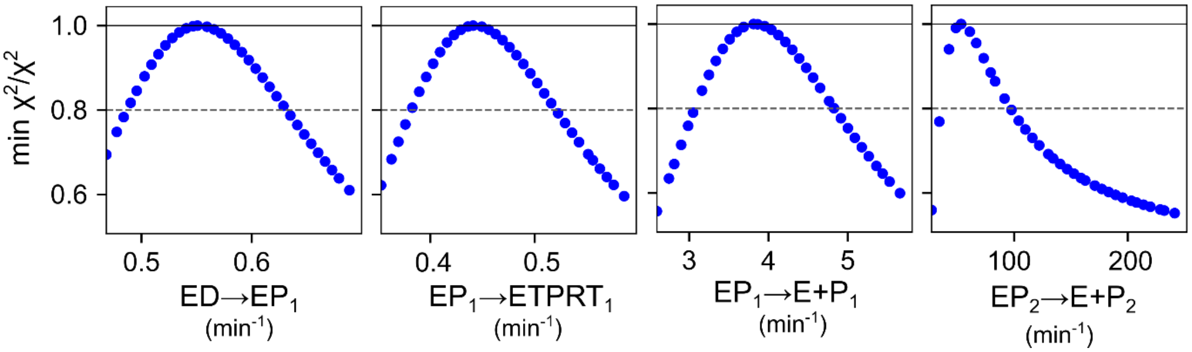


**<u>Figure S3: Confidence Contours for Kinetics of Reaction with CM RNA and Trap DNA.</u>** Confidence contours were generated in KinTek Explorer by fitting to the scheme in Figure 5 (described in the methods section). The dashed line represents the chi$^2$ threshold (0.8) corresponding to the 95% confidence interval for estimated parameters. Best fit parameters and confidence intervals are given in Table S3.

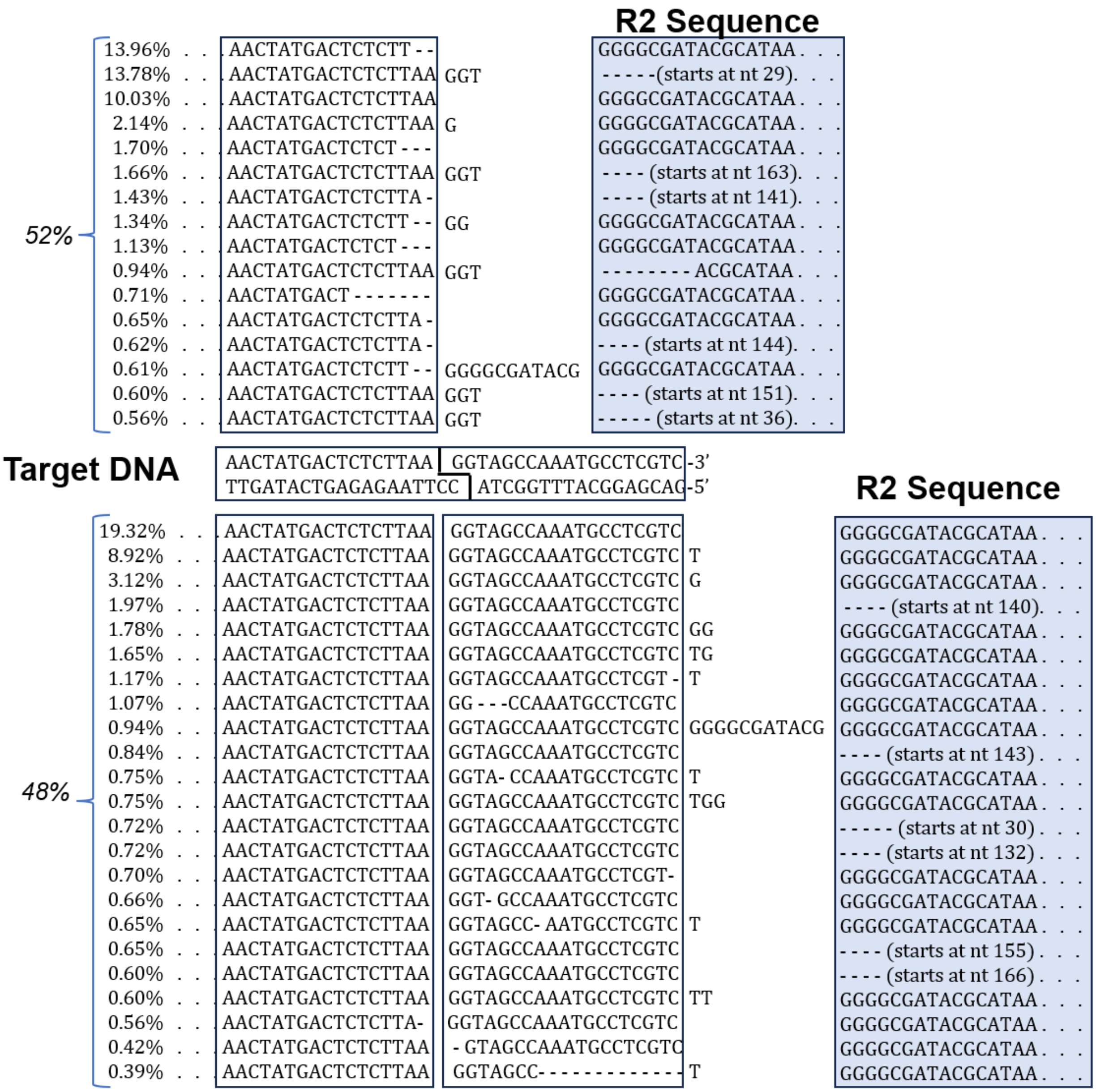


**Figure S4: 5' Junctions Revealed by Next Generation Sequencing.** Amplicon sequencing was performed on a 5' junction targeted PCR product and variability analysis was performed. The various junctions revealed in the analysis are shown, along with percentages of reads corresponding to each resolved junction. Approximately half of the junctions contained sequences starting around the 2nd strand cut site (above target DNA sequence) while the remaining half of the junctions contained sequences where after 2nd strand cleavage, the enzyme polymerized to the end of the DNA template before engaging the end of the cDNA/RNA R2 sequences.

| | | | |
|---|---|---|---|
| Instrument: | MS-IALTQ-06 | Operator ID: | 3548 |
| Acquired: | 3/8/2024 11:31 AM | Reviewed: | 3/8/2024 12:24 PM |

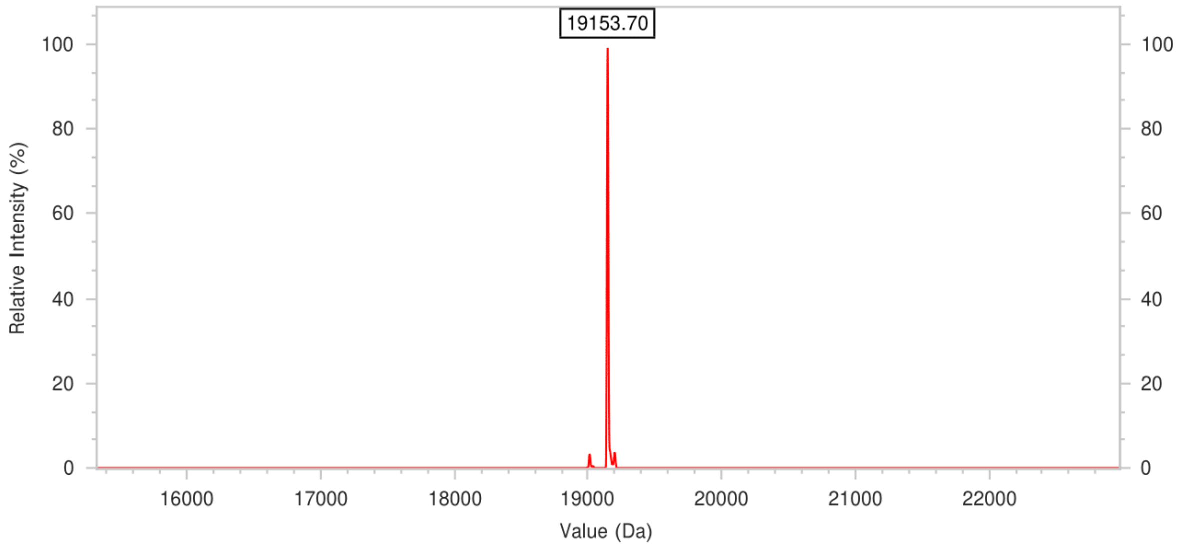


| | |
|---|---|
| **Sequence Name:** | **R2Bm FAM Top 60nt - 3'P** |
| **Sequence:** | **5'- /56-FAM/CAA GCG CGG GTA AAC GGC GGG AGT AAC TAT GAC TCT CTT AAG GTA GCC AAA TGC CTC GTC /3Phos/ -3'** |
| **Calculated Molecular Weight:** | **19151.5** |
| **Measured Molecular Weight:** | **19153.70** |

**Figure S5: Top Strand Oligo Quality Control by ESI Mass Spec**. Quality control data was provided by IDT, the manufacturer of the top strand oligonucleotide used in this study. The oligonucleotide was analyzed by electrospray ionization mass spectrometry and showed a major peak being the full-length oligonucleotide.

A)

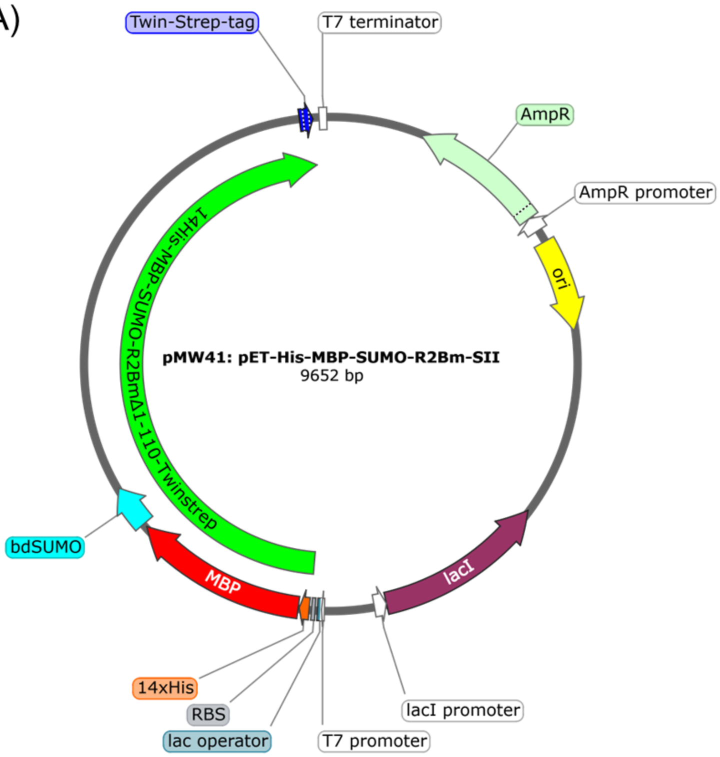


**Figure S6: Map of Plasmid for R2Bm expression.** This plasmid was prepared by Wilkinson et al and is available through Addgene. Locations of the 14xHis, MBP, bdSUMO, and twinstrep tags are shown in orange, red, cyan, and blue, respectively.

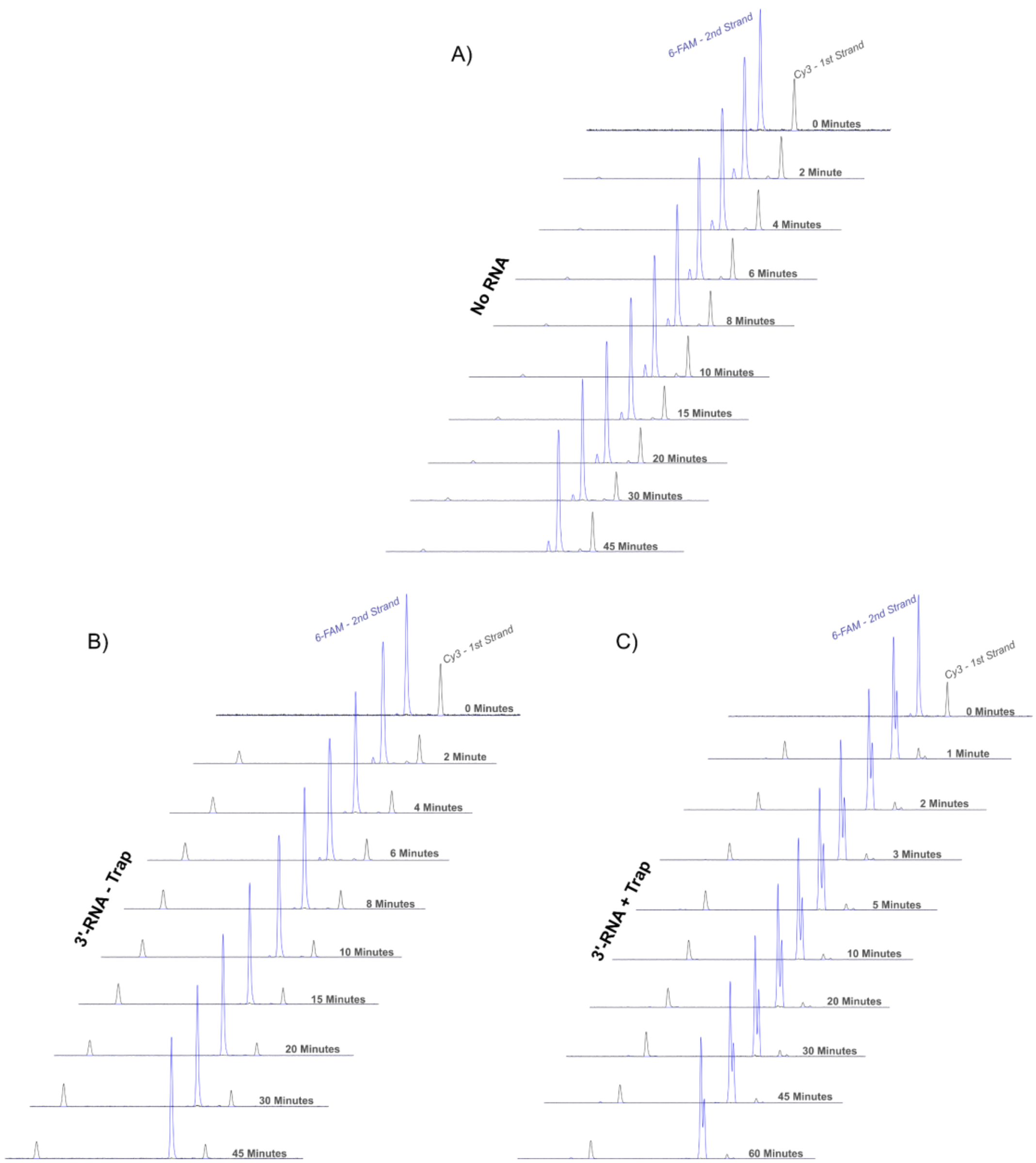


**Figure S7: Electropherograms for Data in Figure 2.** Representative capillary electrophoresis electropherograms from the experiments in panels A and B in Figure 2 are shown.

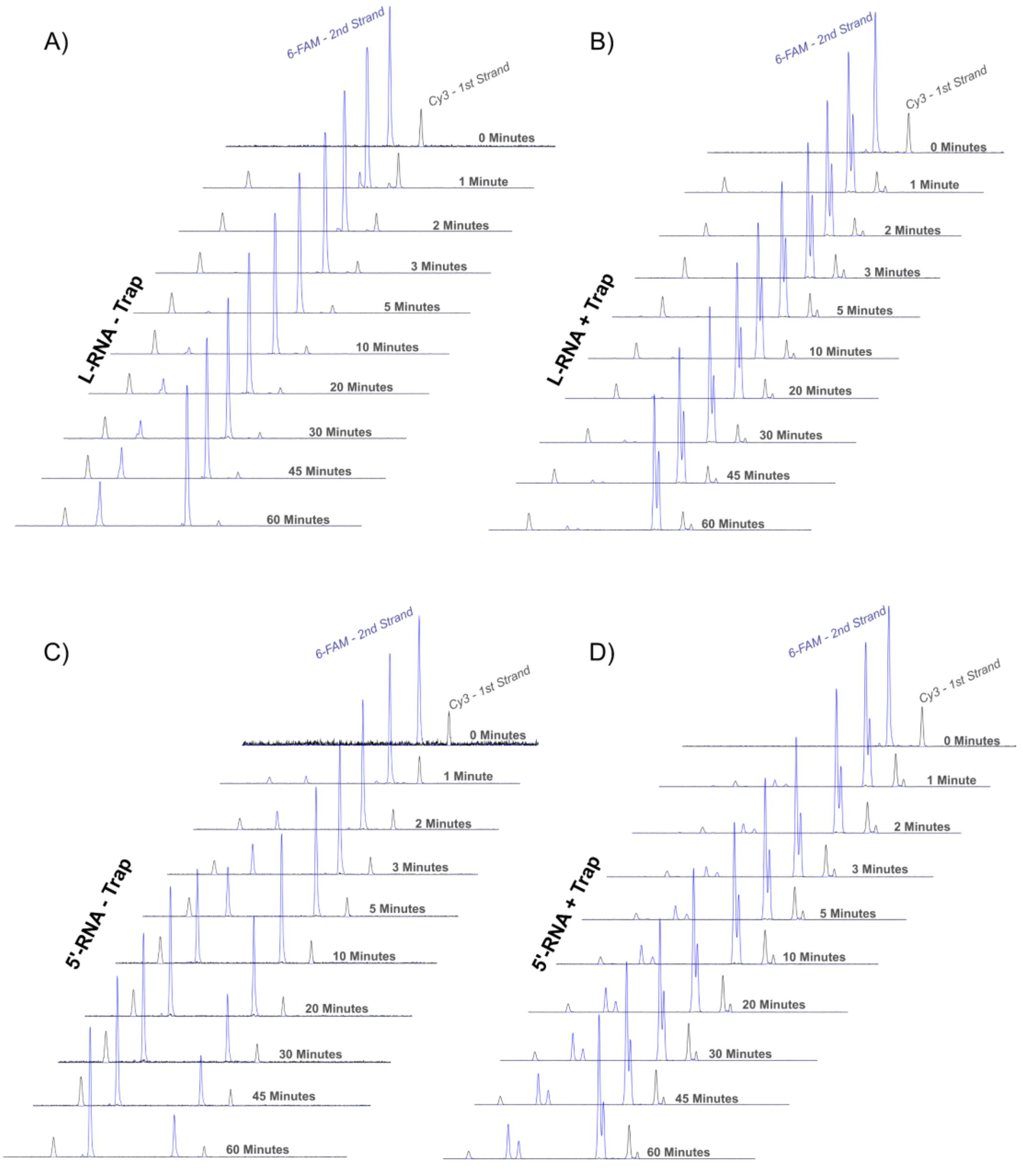


**Figure S8: Electropherograms for Data in Figure 3.** Representative capillary electrophoresis electropherograms from the experiments in Figure 3 are shown.

**Table S1: Rate Constants for First Strand Cleavage and TPRT Kinetics. Related to Figure 4.**

| Step | Best Fit | 95% Confidence Interval |
|---|---|---|
| E + D → ED | 1000 $\mu M^{-1}$ $min^{-1}$ | - |
| E + D → EDx | 1000 $\mu M^{-1}$ $min^{-1}$ | - |
| ED → E + D | 0.1 $min^{-1}$ | - |
| EDx → E + D | 0.043 $min^{-1}$ | 0.038 - 0.049 |
| ED → EP1 | 0.48 $min^{-1}$ | 0.45 – 0.53 |
| EP1 →….→ ETPRT1 ** | 1.58 $min^{-1}$ | 1.26 – 2.05 |
| EP1 → E + P1 | 12.0 $min^{-1}$ | 9.2 – 16.2 |
| * Rate constants in red were locked in the fitting<br>** This was simulated as 10 steps. The rate constant reported here is 1/10$^{th}$ of the value for a single step | | |

**Table S2: Rate Constants for Kinetics with CM-RNA. Related to Figure 5.**

| Step | Best Fit | 95% Confidence Interval |
|---|---|---|
| E + D → ED | 1000 $\mu M^{-1}$ $min^{-1}$ | - |
| E + D → EDx | 245 $\mu M^{-1}$ $min^{-1}$ | 158 – 385 |
| ED → E + D | 0.1 $min^{-1}$ | - |
| EDx → E + D | 0.009 $min^{-1}$ | < 0.024 |
| ED → EP1 | 0.17 $min^{-1}$ | 0.14 – 0.21 |
| EP1 →….→ ETPRT1 ** | 0.40 $min^{-1}$ | 3.2 – 5.3 |
| ETPRT1 → EP2 | 0.54 $min^{-1}$ | > 0.18 |
| EP2 → ETPRT2 | 10 $min^{-1}$ | - |
| EP1 → E + P1 | 2.4 $min^{-1}$ | 1.3 – 3.3 |
| EP2 → E + P2 | 40 $min^{-1}$ | 28 – 67 |
| * Rate constants in red were locked in the fitting<br>** This was simulated as 10 steps. The rate constant reported here is 1/10$^{th}$ of the value for a single step | | |

**Table S3: Rate Constants for Kinetics with CM-RNA with Trap. Related to Figure 5.**

| Step | Best Fit | 95% Confidence Interval |
|---|---|---|
| E + D → ED | 1000 $\mu M^{-1}$ $min^{-1}$ | - |
| E + D → EDx | 250 $\mu M^{-1}$ $min^{-1}$ | - |
| ED → E + D | 0.1 $min^{-1}$ | - |
| EDx → E + D | 0.006 $min^{-1}$ | - |
| ED → EP1 | 0.55 $min^{-1}$ | 0.49 – 0.63 |
| EP1 →….→ ETPRT1 ** | 0.44 $min^{-1}$ | 3.8 – 5.2 |
| ETPRT1 → EP2 | 10 $min^{-1}$ | - |
| EP2 → ETPRT2 | 10 $min^{-1}$ | - |
| EP1 → E + P1 | 3.8 $min^{-1}$ | 3.2 – 4.8 |
| EP2 → E + P2 | 53 $min^{-1}$ | 43 – 92 |
| * Rate constants in red were locked in the fitting<br>** This was simulated as 10 steps. The rate constant reported here is 1/10$^{th}$ of the value for a single step | | |
